# Divergent phenotypic and functional roles of human T follicular helper cells from infancy to adulthood

**DOI:** 10.64898/2026.02.06.704464

**Authors:** Suhas Sureshchandra, Jenna M Kastenschmidt, Erika M Joloya, Zachary W Wagoner, Arjun K Nair, Samuel Kim, Natalie Zane, Gargee Bhattacharya, Alyssa M Monterroso, Evien Cheng, Andrew M Sorn, Mahina Tabassum Mitul, Haven Beares, Germaine Mendez, Timothy B Yates, Fan Zhou, Allyssa Daugherty, Chandrani Thakur, Karl A Brokstad, Douglas Trask, Gurpreet Ahuja, Qiu Zhong, Naresha Saligrama, Rebecca J Cox, Lisa E Wagar

## Abstract

Antibody responses to T-dependent antigens are suboptimal in young children, yet the evolution of T follicular helper cell (Tfh) function across the human lifespan remains poorly defined. Using human tonsils, a physiologically relevant and abundant source of Tfh, we investigated age-associated differences in their repertoire and functional programs. Pediatric tonsils were enriched for cytokine-expressing Tfh subsets with increased clonal diversity and phenotypic plasticity. However, in response to influenza antigens, they exhibited reduced Th1 polarization, diminished IL-21 production, and limited B cell help. Across ages, high neutralizing flu antibody responses were associated with robust Tfh1 activation, which was ICOS dependent in adults but not in children. Interestingly, Tfh depletion strategies revealed enhanced Tfh differentiation from distinct precursors in pediatric donors, yet antibody responses during early life were less reliant on Tfh help. Together, these findings define developmentally programmed differences in Tfh differentiation and function with implications for pediatric vaccine design.

## INTRODUCTION

A critical vulnerability in early life immunity is the inability of young children to generate robust T and B cell responses, resulting in weak, short-lived antibodies that leave them susceptible to infections readily controlled in adults (1). However, it remains unclear whether this deficit stems from intrinsic defects in the quality of T cell help to B cells, from a lack of immunological memory, or both. Most protective antibody responses rely on germinal centers (GCs), where T follicular helper (Tfh) cells secrete essential cytokines, such as IL-21, and express costimulatory molecules, including CD40L and ICOS, that drive B cell selection and long-lived immunity (2,3). In animal models, limiting Tfh function inhibits B cell responses, suppresses GC persistence and function, and reduces plasmablast differentiation (4–6). Despite their central role, the mechanisms that establish and maintain Tfh function across the human lifespan remain poorly defined. Recent work examining circulating Tfh (cTfh) reports marked age-dependent remodeling (7). However, these studies were largely limited to blood and fundamental questions remain regarding human Tfh plasticity within secondary lymphoid tissues, sites where Tfh function. Specifically, it remains unclear how early life Tfh respond to vaccination, to what extent they mediate B cell selection, and whether their helper capacity is inherently distinct from adults.

Since their initial discovery and identification of their transcriptional regulation (8–11), our perspective on Tfh has changed from a discrete cell fate opposing Th1 (12–14), Th2 (15), and Th17 (16,17) fates to a more plastic population of cells capable of acquiring elements of these other programs, depending on antigen type, inflammatory milieu, and tissue context (18). Understanding how such heterogeneity develops during childhood is important because early-life immunity poses unique challenges. Infants experience disproportionate morbidity from respiratory viral infections (19) and often generate weak or short-lived vaccine responses (20–22). However, it remains unclear how Tfh in lymphoid tissues differ in early-life from their adult counterparts in terms of their transcriptional and clonal diversity, differentiation potential, and helper function. Such differences could fundamentally underlie the poor GC quality and suboptimal vaccine responses characteristic of early life.

A key barrier to studying early-life GCs is a lack of functional human systems to model Tfh-B cell interactions. Blood-based analyses offer only a partial picture, as cTfh are an incomplete and temporally restricted snapshot of GC activity (23). Recent studies highlight that early life immunity is highly tissue-localized, with effector activity largely restricted to mucosal sites rather than in circulation (24). Overall, Tfh analysis restricted to peripheral blood may fundamentally miss essential cellular and microenvironmental signals that shape adaptive immunity in children. Fine-needle aspirates of lymph nodes have provided critical insights into vaccine-induced Tfh activity (25) but such sampling is not practical in infants and children. As a complementary approach, we investigated human tonsil Tfh across the age span. Tonsils are one of the few GC-rich mucosal and lymphoid tissues that are accessible from both healthy children and adults (26,27). Furthermore, *in vitro* culture manipulations in tonsil organoids (23,28–30) provide a uniquely suitable platform for dissecting age-dependent Tfh biology and ontogeny in a tissue context.

Influenza vaccination presents a useful model for dissecting Tfh function in early life and enables us to separate the influence of prior antigen exposure from age-dependent, cell-intrinsic differences in Tfh differentiation and function (31,32). In adults, influenza vaccines can induce Tfh1 (25) and IL-21 production (33), which correlate with neutralizing antibodies. Live-attenuated influenza vaccines (LAIVs) induce Tfh activation in tonsils as early as 2-3 days post-vaccination (34). However, infants broadly mount poor GC and humoral responses to influenza vaccines (35). In this work, we sought to understand whether these suboptimal antibody responses in children reflect intrinsic Tfh limitations, either in their differentiation or function, in early life. Using pediatric and adult tonsils, including from recently LAIV-immunized donors, we mechanistically dissected the relative contributions of Tfh to antibody responses and how these contributions change during childhood.

## RESULTS

### Clonally related resting and cytokine-expressing Tfh subsets are enriched in pediatric tonsils

We began our analysis by first investigating Tfh heterogeneity in human tonsils from pediatric and adult donors (**Table S1**) using an unbiased transcriptomic approach (**Fig. 1A**). Single-cell RNA-seq (scRNA-seq) with paired cell-surface protein and TCR analysis of CD4 T cells from pediatric (2-3 years old) and adult donors (24-39 years old, n=6/group) revealed considerable inter-group Tfh heterogeneity (**Fig. 1B and fig S1A)**, defined both by gene expression (**Fig. 1C and fig S1B)** and protein markers (**Fig. 1D and fig S1C)**. We identified the expected Tfh states, including germinal center Tfh (GC-Tfh, expressing high *CXCR5*, *PDCD1*, *BCL6*, *CD57*) and intermediate *BCL6* expressing Tfh (36). We also identified two novel Tfh states characterized by distinct cytokine expression profiles. One of these expressed high *IL10* mRNA (termed Tfh-IL10 hereafter), and another had high expression of *TNF* mRNA (Tfh-TNF) (**Fig. 1C**). As expected, non-Tfh memory subsets such as Th1 and central memory (TCM) cells were elevated in adults **(fig S1D)**, but all four Tfh clusters were enriched in pediatric donors (26) (**Fig. 1E**).

**Fig. 1:**
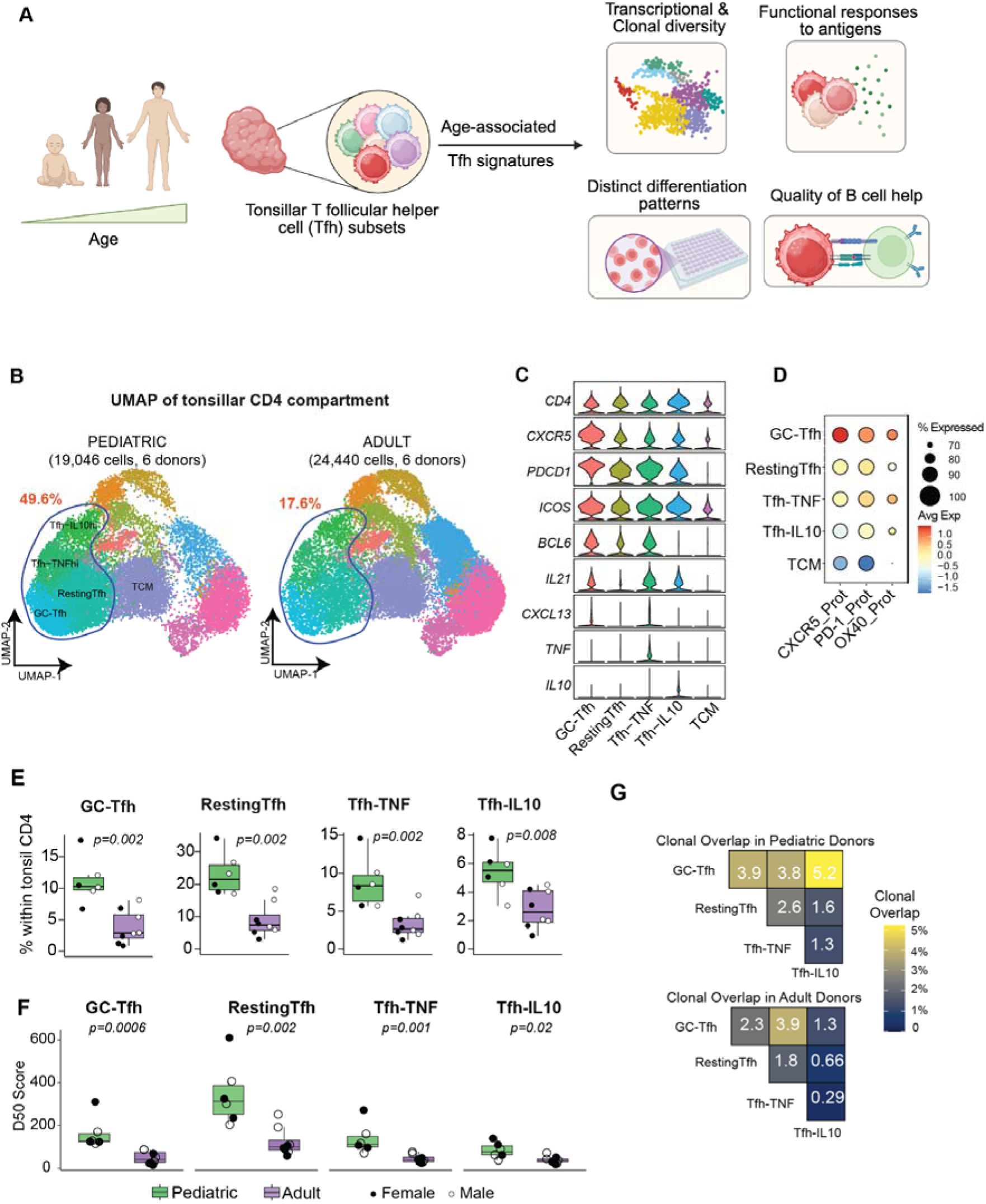
Early life transcriptional heterogeneity and clonal characteristics of tonsillar T follicular helper cells. (A) Experimental Design. Tonsil lymphocytes were isolated from healthy pediatric and adult donors. Cryopreserved cells were revived, and Tfh compartments were compared between the two groups using single-cell genomics, flow cytometry, and functional assays. (B) UMAP of 48295 CD4 T cells from palatine tonsils from 12 donors (n=6/group). Tfh clusters are highlighted with their total percentages (% of CD4) compared. (C) Violin plots comparing mRNA expression of key Tfh and cytokine markers expressed by diverse Tfh subsets in tonsils. (D) Bubble plot comparing surface protein expression of key Tfh markers among diverse Tfh subsets in the tonsils, relative to resting central memory CD4. The size of the bubble indicates the proportion of cells within the cluster expressing the marker, and the color indicates the magnitude of expression. (E) Boxplots comparing proportions of Tfh subsets within the total CD4 compartment in tonsils from pediatric (<5 years old, n=6) and adult donors (n=6). (F) Boxplots comparing clone diversity (D50) of Tfh subsets between pediatric and adult donors. D50 is a diversity estimate representing the minimum number of distinct clonotypes amounting to greater than or equal to 50% of the total TCRs counted in the sample. Boxplots show the median, with hinges indicating the first and third quartiles and whiskers indicating the highest and lowest values within 1.5 the interquartile range. (G) Heatmap comparing clonal overlap among Tfh subsets in pediatric (top) and adult (bottom) donors. Clonal overlap was calculated using immunarch.

We previously demonstrated that Tfh represent the most clonally expanded T cell subset in pediatric tonsils, even more so than CD8 T cell subsets in the same donors (23). However, pediatric Tfh subsets still maintained greater TCR diversity than their adult counterparts, suggesting accumulation of dominant clones over time (**Fig. 1F**). We next assessed the extent of clonal overlap between Tfh subsets (**Fig. 1F**). Resting Tfh were the most clonally diverse among the four Tfh subsets (**Fig. 1F**) and exhibited the least clonal sharing with other Tfh subsets (**Fig. 1G**). In contrast, the cytokine-expressing Tfh were the most clonal and exhibited substantial clonal overlap with GC-Tfh (**Fig. 1G**). However, the extent of sharing was significantly higher in pediatric donors (**Fig. 1G and fig S1E)**, suggesting greater phenotypic and clonal plasticity within Tfh during early life.

### Pediatric Tfh display reduced functional capacity for B cell help and cytokine production

Given the differences in overall frequencies and clonality of Tfh subsets, we next asked if Tfh were functionally different between the two age groups. We began by comparing transcriptional profiles of the GC-Tfh subset using pseudobulk differential analysis. GC-Tfh from adult donors had upregulated expression of key immune and cytokine signaling molecules, including master regulators of Th1 responses (*STAT1* and *STAT4*), cytokine modulators (*CD96* and *SOCS1*) and mediators facilitating B cell help (*CD84*, encoding SLAMF5) (**Fig. 2A and fig S2A)**. Flow cytometry analysis confirmed higher surface expression of B cell help markers (OX40, CD40L, CD84) at the protein level on adult relative to pediatric GC-Tfh (**Fig. 2B**).

**Fig 2:**
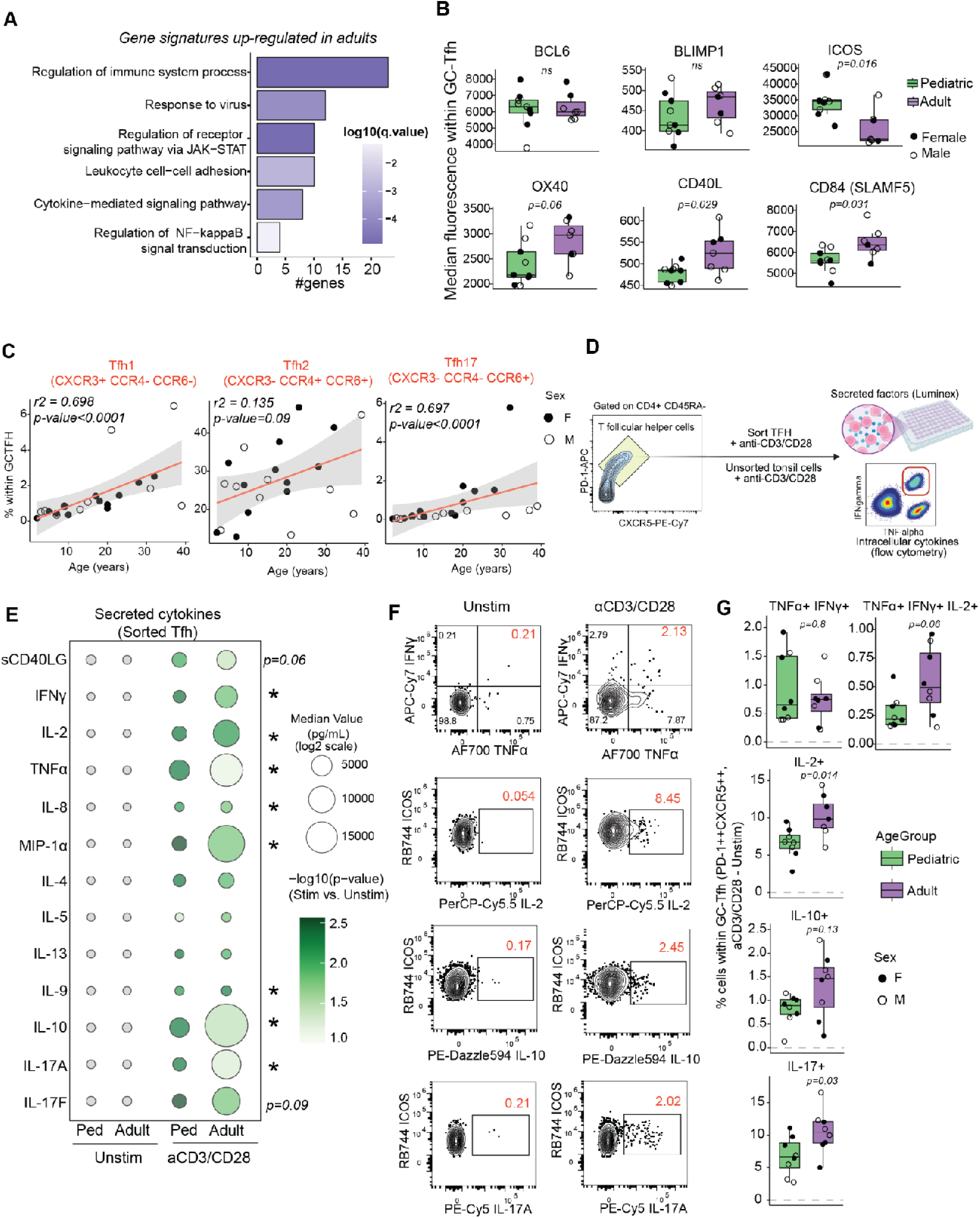
Functional Tfh responses are attenuated during early life. (A) Bar graph highlighting functional enrichment of gene signatures up-regulated in adult Tfh compared to their pediatric counterparts. The X-axis represents the number of genes in each Gene Ontology (GO) term, and the color represents statistical significance. (B) Boxplots comparing protein expression (median fluorescence or %, as highlighted) of key surface protein and transcription factors within Tfh (CXCR5++PD-1++) in pediatric and adult donors, measured using flow cytometry. (C) Dot plots highlighting the proportion of Tfh1, Tfh2, and Tfh17 cells, defined by chemokine receptor expression, within *ex vivo* Tfh. The X-axis represents the age of the donor, and the Y-axis represents the proportion of cells as a % of Tfh. P-values were calculated using two-sided Spearman’s rank correlation. (D) Experimental Design for cytokine assays. Tonsillar cells were stimulated overnight with anti-CD3/CD28, and cytokine responses relative to unstimulated controls were measured using intracellular cytokine staining and flow cytometry. Additionally, Tfh were FACS-sorted and stimulated with anti-CD3/CD28 for 16 hours, and secreted cytokines in the supernatants were measured using Luminex. (E) Bubble plot comparing key cytokines from conditioned media from sorted Tfh stimulated (anti-CD3/CD28) or unstimulated (-) overnight between pediatric and adult donors. Secreted cytokines were measured using Luminex. For each analyte, average values across pediatric and adult donors were calculated. The bubble size denotes the average pg/mL of the cytokine, and the color intensity indicates statistical significance (-log10(p-value)) for comparisons between stimulated conditions in pediatric and adult donors. Deeper purple denotes higher statistical significance calculated using paired Wilcoxon signed-rank tests. (F) Representative gating strategy for evaluation of cytokine-expressing Tfh subsets. All gates were drawn relative to fluorescence-minus-one controls. (G) Box plots comparing proportions of cytokine-expressing cells within CXCR5++PD-1++ Tfh in anti-CD3/CD28 and unstimulated tonsil cells. The y-axis represents % values, corrected for background in unstimulated conditions. Boxplots show the median, with hinges indicating the first and third quartiles and whiskers indicating the highest and lowest values within 1.5 the interquartile range.

Interestingly, we observed no age-related differences in protein expression of the Tfh master regulator BCL6 nor its antagonist transcription factor BLIMP1 in GC-Tfh (**Fig. 2B and fig S2B)** (9), suggesting limited qualitative differences in enforcement of the Tfh program related to age. Since Tfh have been shown to incorporate elements of other Th differentiation programs (14,26,37), we next asked whether early-life Tfh preferentially skew towards a particular Th program over others. Using chemokine receptor expression (CCR4, CCR6, and CXCR3), we quantified Tfh1 (CXCR3+CCR4-CCR6-), Tfh2 (CXCR3-CCR4+CCR6-), and Tfh17 (CXCR3-CCR4+CCR6+) subsets within GC-Tfh (38) **(fig S2C)**. Our analysis showed that GC-Tfh were preferentially of a Tfh2 flavor in both pediatric and adult tonsils (**Fig. 2C**). However, adult GC-Tfh were better able to acquire Th1 and Th17 phenotypes and modestly higher Th2 phenotypes compared to their pediatric counterparts (**Fig. 2C**).

We next tested if age-dependent Tfh differences translated tino altered cytokine responses following TCR engagement. Sorted tonsillar GC-Tfh (CXCR5++PD-1++) cells or total tonsillar mononuclear cells were stimulated with anti-CD3/CD28 overnight and secreted and intracellular cytokines were quantified (**Fig. 2D**). Consistent with the Th bias data, GC-Tfh from adults secreted higher levels of Th1 (IFNL, TNFL), Th17 (IL-17A), and pleiotropic cytokines such as IL-10 (**Fig. 2E**). We also observed higher frequencies of polyfunctional (IL-2, TNFL, IFNL-producing) and IL-17-producing Tfh in adults (**Fig. 2F and 2G**). This was surprising, given that both Tfh-IL10 and Tfh-TNF subsets were abundant in pediatric donors (**Fig. 1E**), and suggested post-transcriptional regulation or differences in effector maturity between adult and pediatric Tfh. Collectively, these findings show that the proportion of functional Tfh cells is not diminished in early life but the magnitude of their response is reduced compared to adults.

### Age-dependent Tfh responses to influenza vaccination *in vivo*

We next investigated how Tfh respond to influenza vaccination *in vivo* between the two age groups. We analyzed tonsil samples from a cohort of healthy donors in Bergen, Norway, who received tonsillectomies 5-9 days after intranasal LAIV (**Fig. 3A**) and compared their *ev vivo* Tfh phenotypes with a cohort of donors who had not been recently vaccinated (indicated by “No recent vax”). Pediatric donors exhibited significant expansion of CXCL13-expressing GC-Tfh following vaccination (**Fig. 3B-C**) and showed greater upregulation of ICOS and interferon-stimulated genes compared to adults (**Fig. 3D**). To assess functional cytokine responses, we cultured tonsil cells from vaccinated and unvaccinated donors for 4 days without additional stimulation and measured spontaneously secreted cytokines in the supernatants. In vivo vaccination elicited a stronger Th1 cytokine response in adults compared to pediatric donors (**Fig. 3E**). These findings demonstrate distinct age-dependent Tfh activation programs in response to *in vivo* influenza vaccination in the tonsil.

**Fig 3:**
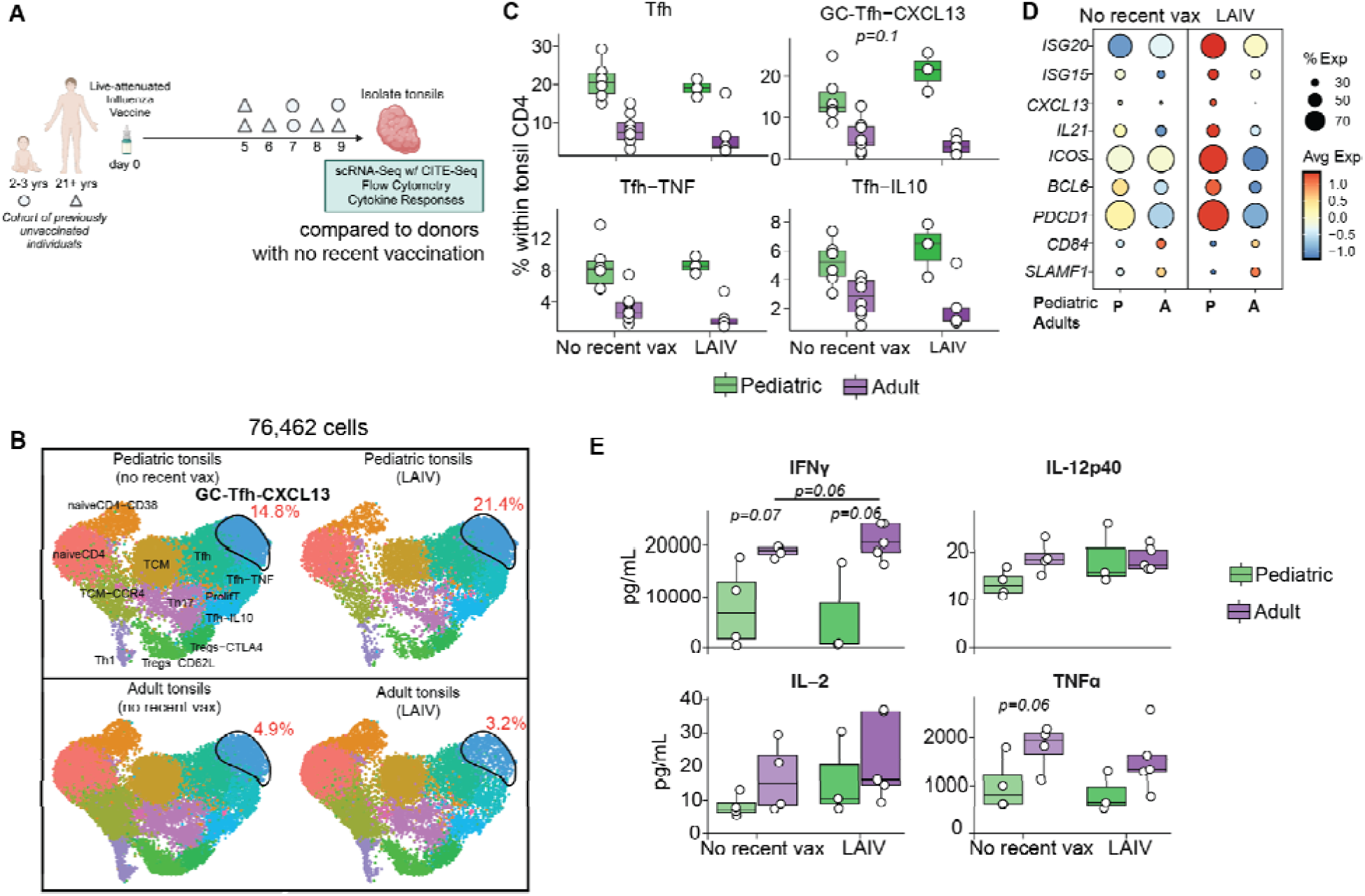
Tfh responses to *in vivo* influenza vaccination. (A) Experimental design. (B) UMAP of 76,462 CD4 T cells from palatine tonsils from recently vaccinated and unvaccinated pediatric (top) and adult donors (bottom). Mature CXCL13-expressing GC-Tfh clusters are highlighted with their total percentages (% of CD4) compared across groups. (C) Boxplots comparing proportions of Tfh subsets within the total CD4 compartment in the tonsils from vaccinated and unvaccinated pediatric (<5 years old) and adult donors. P-values highlight comparison between no recent vax and LAIV group using unpaired t-test within each age group. (D) Bubble plots comparing mRNA levels of key Tfh-associated markers within the GC-Tfh cluster between unvaccinated and vaccinated donors. The color of the bubble denotes the normalized transcript counts, and the size of the bubble indicates % of cells within the cluster that express the corresponding marker. (E) Boxplots comparing secreted levels of Th1 cytokines in supernatants from unstimulated organoids generated from vaccinated and unvaccinated donors. Labels on the x-axis indicate in vivo vaccination status. Group differences were compared using unpaired Mann-Whitney U tests (two-sided). Boxplots show the median, with hinges indicating the first and third quartiles and whiskers indicating the highest and lowest values within 1.5 the interquartile range.

### Tonsil immune organoids reveal age-dependent differences in Tfh responses to influenza antigens

While *in vivo* vaccination data revealed minor age-dependent differences in Tfh responses, there are several caveats limiting the interpretation of this data. Sampling from a single time point cannot capture Tfh dynamics and post-vaccination cell trafficking complicates interpretation of immune responses. To complement *in vivo* studies and enable mechanistic investigation of age-dependent Tfh responses to vaccine antigens, we used tonsil immune organoids to longitudinally track Tfh dynamics over two weeks post-vaccination (**Fig. 4A**). Cryopreserved cells from healthy pediatric and adult donors were cultured as organoids as previously described (28,39) and T cell responses to LAIV or wild-type influenza virus were assessed using scRNA-seq paired with TCR, flow cytometry, and cytokine assays (**Fig. 4A**).

**Fig 4:**
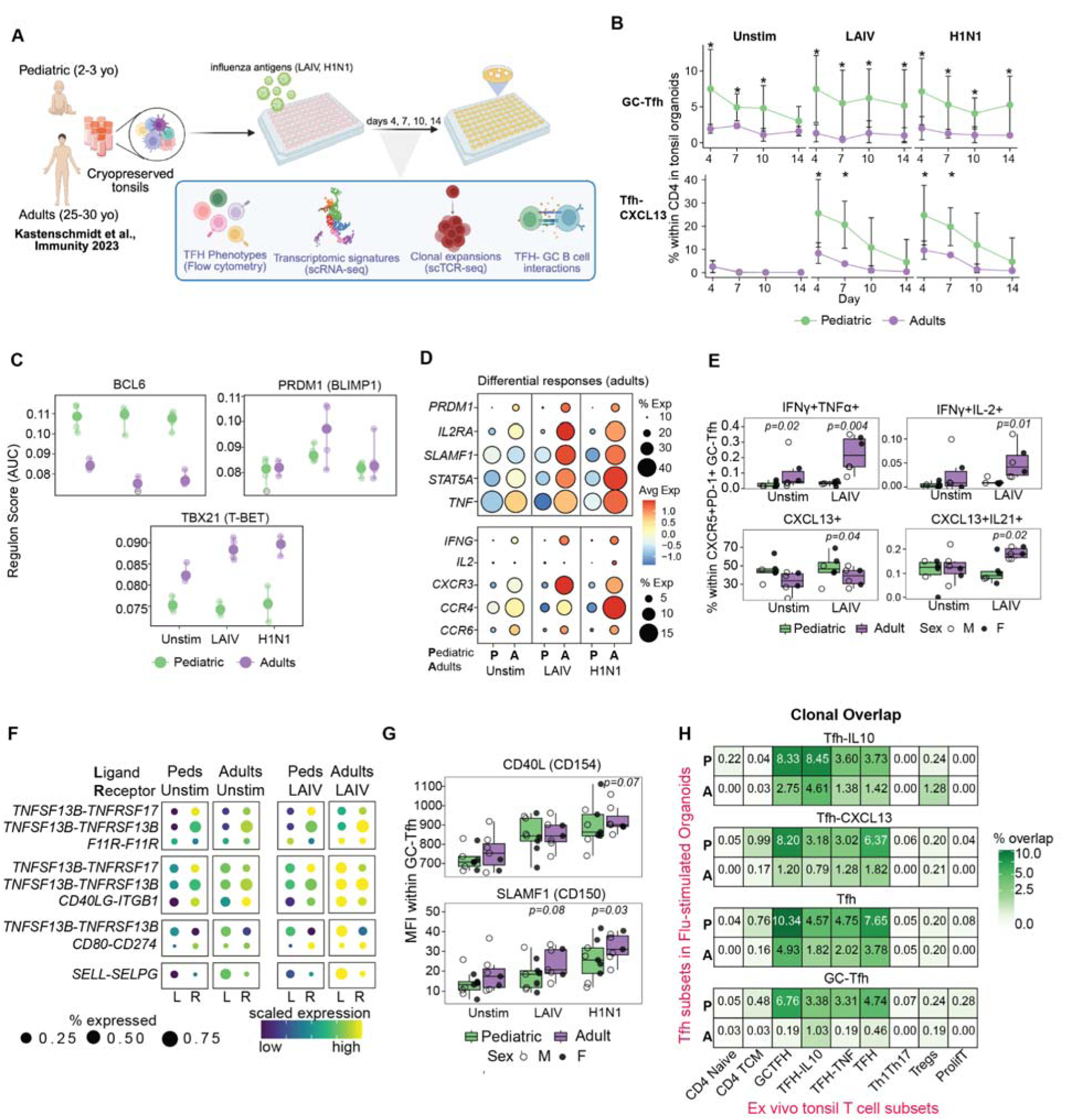
Divergent T cell responses to influenza antigens in pediatric and adult donors. (A) Experimental design for measuring Tfh responses to influenza in tonsil immune organoids. Organoids from pediatric and adult tonsils were generated, and Tfh readouts were measured longitudinally on days 4, 7, 10, and 14 post stimulation with either live-attenuated influenza vaccine (LAIV) or wild-type virus. scRNA-seq was performed on the total CD4 T cell compartment, with orthogonal TCR and cell surface protein measurements. (B) Line graphs comparing proportions of key Tfh subsets in immune organoids over time (x-axis) within the total CD4 compartment (y-axis). At each time point, group differences were compared using unpaired Mann Whitney U test (* - p<0.05) (C) Dot plot comparing regulon AUC of targets of key transcription factors in mature GCTfh-CXCL13 subset on day 7 in unstimulated and flu-stimulated organoids from pediatric and adult donors. Cell level TF activity scores were aggregated by calculating mean AUC per donor per stim condition. (D) Bubble plots comparing mRNA levels of differentially expressed genes in the GCTfh-CXCL13 subset between unstimulated and flu-stimulated conditions. Gene signatures up-regulated solely adult are shown. The color of the bubble denotes the normalized transcript counts, and the size of the bubble indicates % of cells within the cluster that express the corresponding marker. (E) Box plots comparing proportions of cytokine-expressing cells within CXCR5++PD-1++ Tfh on day 7 in LAIV and unstimulated tonsil organoids. Boxplots show the median, with hinges indicating the first and third quartiles and whiskers indicating the highest and lowest values within 1.5 the interquartile range. Within each stim condition, group differences were compared using unpaired Mann Whitney U test. (F) Bubble plot highlighting the strength of ligand-receptor interactions predicted using NicheNet. Statistically significant interactions were identified from pseudobulk differential analysis performed for ligands in sender cells and receptor genes in receiver cells (minimum 10 cells, p < 0.05; log2FC > 0.5) (G) Box plots comparing surface expression of key markers associated with B cell help in immune organoids. Boxplots show the median, with hinges indicating the first and third quartiles and whiskers indicating the highest and lowest values within 1.5 the interquartile range. Within each stim condition, group differences were compared using unpaired Mann Whitney U test. (H) Heatmap comparing clonal overlap of Tfh subsets in flu-stimulated organoids on days 4, 7, 10, and 14 combined (y-axis) and ex vivo CD4 subsets (day 0, x-axis) in pediatric (top) and adult (bottom) donors. Clonal overlap was calculated using immunarch.

Single-cell analyses of roughly 400,000 T cells revealed expected heterogeneity as previously described (28), including naive, memory, Tfh subsets, Th1, Th17, regulatory T cells (Tregs), activated, and proliferating populations **(figs S3A and S3B)**. Longitudinal analysis of T cell cluster frequencies under different stimulation conditions highlighted striking age-dependent differences in Tfh subsets (**Fig. 4B**). While resting GC-Tfh frequencies remained stable across all conditions, influenza antigen stimulation expanded cytokine-expressing Tfh subsets. Specifically, pediatric organoids showed early expansion of *CXCL13* expressing Tfh, as observed during *in vivo* vaccination, and late expansion of Tfh-IL10, both at significantly higher levels than adults (**Figs. 4B and S3C)**. In contrast, responses in adult organoids were dominated by Th1 polarization, with expansion of CXCR3+ Th1 and proliferating T cells between days 7 and 10, and a peak in antigen-responsive OX40+ T cells on day 7 - all at higher magnitudes than pediatric donors **(fig S3C)**. Together, these findings demonstrate qualitatively distinct responses to flu antigens, suggesting age-dependent programming of helper versus effector T cell fates.

### Poor Th1, but enhanced Tfh differentiation in response to influenza vaccine in early life

Although GC-Tfh frequencies remained unchanged with influenza stimulation, differential analysis revealed age-specific activation states in response to virus or vaccine (**Figs. 4C and 4D**). Within GC-Tfh, targets of BCL6 were downregulated while those of BLIMP-1 (encoded by *PRDM1*) were upregulated with influenza antigen stimulation in adults - a pattern absent in pediatric donors (**Fig. 4C**). To our surprise, type-I interferon responses were strongest in pediatric Tfh (**fig S3D**), reflecting a heightened antiviral state. On the other hand, antigen-specific effector responses, including upregulation of activation markers (*IL2RA*/CD25) (**Fig. 4D**), effector cytokines (*IFNG*, *TNF*) (**Figs. 4D and 4E**), T-bet (encoded by *TBX21*) signaling targets (**Fig. 4C**), and Th1/Th17 signatures (*CXCR3*, *CCR4*, *CCR6*) (**Fig. 4D**), were prominent in adult Tfh from influenza stimulated organoids. Furthermore, adult organoids secreted significantly higher Th1 and Th17 cytokines in response to influenza antigens (**Figs 4E and S3E)**. While pediatric Tfh maintain a canonical GC program, these data show that adult Tfh better progress towards an effector-like state in response to influenza stimulation.

In concordance with transcriptomic data (**Fig. 4B**), CXCL13 expressing Tfh were more abundant in pediatric donors (**Fig. 4E**). However, functional Tfh cells co-expressing IL-21 and CXCL13 expanded exclusively in adult cultures following influenza stimulation (**Fig. 4E**). This led us to hypothesize that while pediatric Tfh can recruit B cells to GCs, their capacity to provide functional help through IL-21 may be limited. Consistent with this hypothesis, cell-cell interaction analysis using NicheNet revealed stronger receptor-ligand interactions between GC-Tfh and B cell subsets in adults following LAIV stimulation (**Fig. 4F**). Adults exhibited enhanced GC-Tfh interactions with both memory B cells (via BCMA and TACI) and light zone GC B cells (via PD-L1, TACI, CD29) compared to children (**Fig. 4F**). Flow cytometry also confirmed higher induction of two key markers on Tfh critical for B cell help: CD40L (in response to virus stimulation) and SLAMF1 (in response to both vaccine and virus) (**Fig. 4G**). Overall, we conclude that Tfh help to B cells is a limiting factor for GC responses in pediatric donors.

### Clonal analysis reveals age-dependent differences in Tfh ontogeny differentiation

T cell receptor (TCR) sequences can serve as unique barcodes that enable us to track the origin, proliferation, and differentiation potential of a given T cell clone. We used TCR sequence analysis to understand the expansion potential of different CD4 T cell populations, including Tfh, following antigen stimulation. Analysis of clone size distribution in influenza-stimulated organoids revealed no major age-related differences in Tfh clonality nor in other effector CD4 T cell subsets. The only exception was proliferating T cells, which were more clonally restricted in adult-derived organoids following vaccination **(fig S3F)**. Even Th1 cells, which were proportionally lower in pediatric donors, exhibited clone sizes comparable to those of Th1 in adults **(fig S3F)**. To investigate the cellular origins of Tfh subsets following influenza stimulation, we used the TCR to track clonal relationships between *ex vivo* tonsil CD4 T cell subsets and Tfh populations in organoids (on day 7). Clonal analysis between day 0 and day 7 revealed that naive to Tfh differentiation occurred at a higher rate in pediatric organoids (**Fig. 4H**). Interestingly, GC-Tfh showed strikingly reduced clonal sharing with cytokine-expressing Tfh in adult organoids compared to pediatric organoids (**Fig. 4H**), suggesting that either adult GC-Tfh were less plastic following stimulation or that clonally restricted populations were selectively maintained within the GC. Overall, our TCR analysis revealed that while clonal expansion of antigen-responsive tonsillar Tfh is similar in children and adults, their origins and plasticity differ by age.

### Tfh activation predicts antibody responses independent of age

Given the central role of Tfh in supporting antibody affinity maturation, we hypothesized that functional differences we observed in pediatric Tfh might translate to impaired antibody responses. We leveraged a cohort of 100 donors with previously characterized influenza vaccine responses from tonsil organoids (30) and focused our analysis on donors who mounted a robust neutralizing antibody response to LAIV. This strategy enabled a fair comparison of Tfh features between responding pediatric and responding adult donors rather than comparing all children (including non-responders) to adults (**Fig. 5A**). Independent of age, responders showed a peak in CXCR3 expressing Tfh (Tfh1) on day 7 that subsided over time in both groups (**Fig. 5B**). Notably, frequencies of activated (CD38++HLA-DR++) Tfh1 peaked early on day 4 in all responders but displayed distinct kinetics thereafter; activated Tfh1 contracted over time in adults (*p=0.019*) but were sustained throughout the culture period in pediatric donors (*p=0.13)* (**Fig. 5C**).

**Fig 5:**
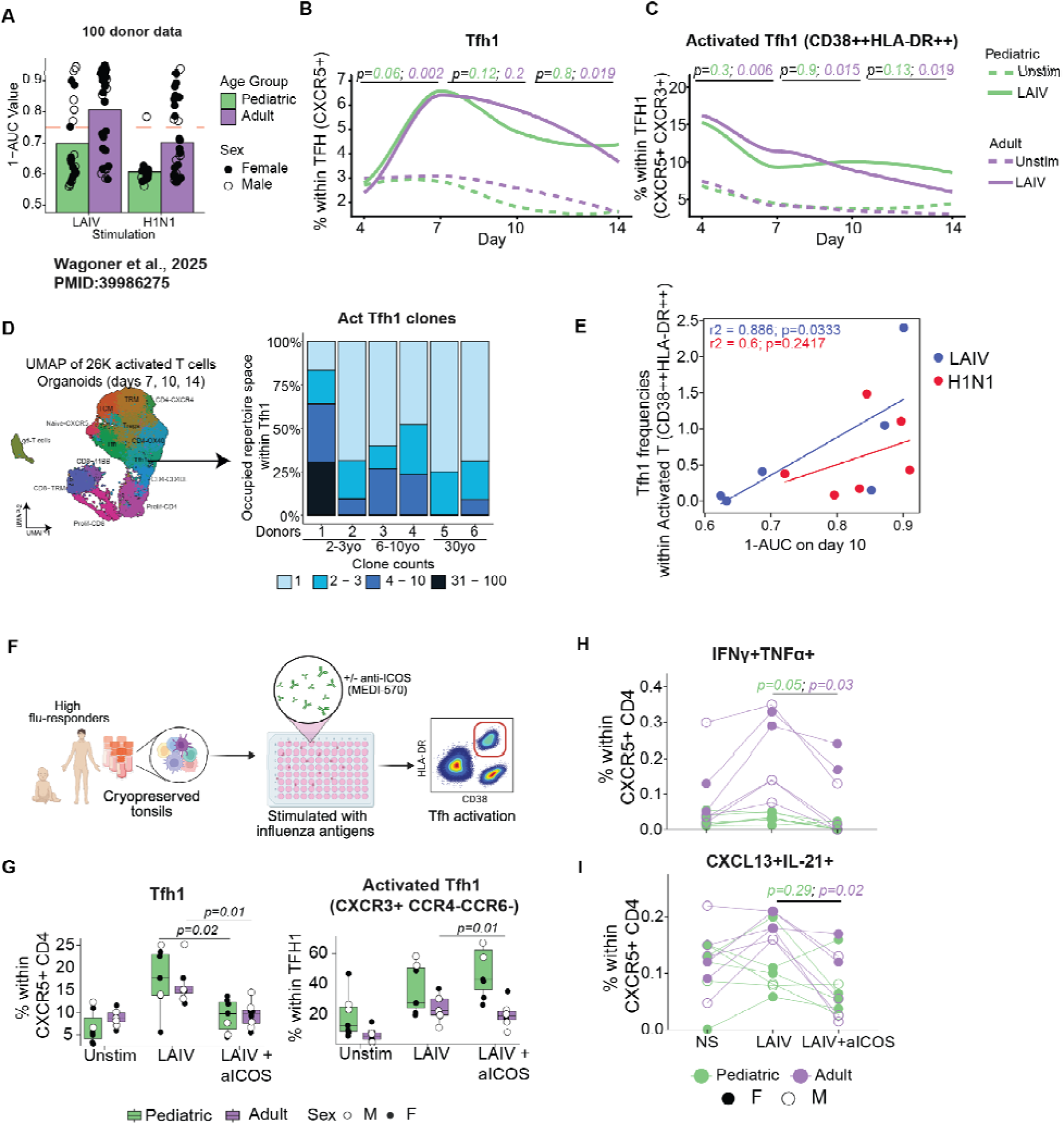
Tfh activation correlate with antibody responses regardless of age. (A) Comparison of neutralization data, quantified by area under the curve (AUC) for organoids stimulated with influenza antigens from pediatric and adult donors. Donors were 1-AUC value greater than 0.75 were categorized as high responders. Original data is described in (30). (B-C) Line graphs comparing (B) CXCR3 expressing Tfh (Tfh1) and (C) HLA-DR++CD38++ expressing Tfh1 in unstimulated (dashed lines) and LAIV stimulated (solid lines) organoids from pediatric and adult responders. At each time point, stim comparisons (LAIV vs. Unstim) were performed within each age group independently (green - pediatric; purple - adult) using paired Wilcoxon signed-rank tests (two-sided). (D) Clone size distribution of antigen-responsive cluster of activated Tfh1. Each bar represents a donor with their age highlighted on x-axis. (E) Dot plot comparing activated Tfh1 frequencies with magnitude of neutralizing antibodies (1-AUC values) on day 10. P-values were calculated using two-sided Spearman’s rank correlation. (F) Experimental design for Tfh functional blocking. Organoids were generated from pediatric and adult donors (n=6/group), stimulated with influenza vaccine, and treated with or without anti-ICOS antibody. Tfh activation was measured using flow cytometry on day 7. (G) Frequencies of CXCR3+ Tfh (Tfh1) as a % of Tfh (left), and activated Tfh1, as a percent of Tfh1 (right). Boxplots show the median, with hinges indicating the first and third quartiles and whiskers indicating the highest and lowest values within 1.5 the interquartile range. Within each age group, stimulation specific differences were tested using paired Wilcoxon signed-rank tests. For age-differences within each stimulation, unpaired Mann-Whitney U tests were performed. (H-I) Dot plots comparing proportions of cytokine-expressing cells within CXCR5+ Tfh on day 7 in unstimulated and LAIV treated organoids with or without ICOS blockade. Within each age group, ICOS specific differences were tested using paired Wilcoxon signed-rank tests (green - pediatric; purple - adult)

To understand whether persistent Tfh1 activation in pediatric donors reflected altered clonal expansion dynamics, tonsil organoids were stimulated with LAIV or influenza virus (A/California 2009 H1N1) and activated T cells (CD3+ HLA-DR+ CD38+) were sorted on days 7, 10, and 14 for single-cell RNA and TCR sequencing to track clonality over time **(fig S4A)**. Activated Tfh1 cells (*CXCR5*+*PDCD1*+*CXCR3*+) exhibited clear antigen-specific expansion in influenza-stimulated organoids **(figs S4B and S4C)**, enabling clonal tracking analysis. Despite differences in their frequencies, Tfh1 displayed comparable clonality across age groups, indicating similar proliferative capacity (**Fig. 5D**). Critically, activated Tfh1 frequencies predicted antibody neutralization capacity independent of age (**Fig. 5E**), establishing activated Tfh1 activation as a functionally relevant correlate of vaccine-induced protection across both pediatric and adult donors.

### Adult Tfh show greater functional dependence on ICOS signaling

Having established that activated Tfh1 cells correlate with antibody responses in both age groups, we tested whether the mechanisms maintaining Tfh activation following vaccination differed with age. Given that ICOS is a key costimulatory pathway for Tfh activation and function (40), we tested whether pediatric and adult Tfh had differential sensitivity to ICOS blockade. Tonsil cells from high-responding donors (n=6 per age group) were cultured as organoids, stimulated with LAIV, and treated with or without anti-ICOS antibody for one week, after which Tfh activation was assessed (**Fig. 5F**).

ICOS blockade altered core Tfh features in an age-dependent manner. In both groups, inhibiting ICOS signaling led to a contraction of PD-1hi, GC-associated Tfh **(fig S4D)**, with a more pronounced effect in adults. Frequencies of CXCR3-expressing Tfh1 were reduced in both groups (**Fig. 5G, left**), indicating that ICOS signaling contributes broadly to Tfh1 polarization. However, a striking age-specific difference emerged when examining Tfh1 activation; while their frequencies were largely unchanged with ICOS inhibition in pediatric donors, activated Tfh1 cells were significantly diminished in adults (**Fig. 5G, right**). Consistent with this, both Th1 (**Fig. 5H**) and IL-21 (**Fig. 5I**) responses were markedly reduced in adults following ICOS blockade, revealing that adult Tfh were more functionally dependent on ICOS signaling for acquiring effector function during a vaccine response (**Figs. 5G-I**). Taken together, these findings indicate that activated Tfh1 predict antibody responses in both age groups. However, the mechanisms sustaining Tfh activation differ based on age, with adult Tfh reliant on ICOS signaling, while pediatric Tfh maintain activation through ICOS-independent pathways.

### Tfh regeneration in immune organoids reveals age-dependent differentiation programs

Given the distinct signals Tfh require to respond to antigens during early life, we sought to understand the events underlying their initial differentiation in young children and adults. To do this, we depleted pre-existing CXCR5+PD-1+ CD4 T cells before organoid culture, then compared T and B cell responses in Tfh-depleted cultures vs. Tfh-reconstituted control organoids (**Figs 6A and 6B**).

**Fig 6:**
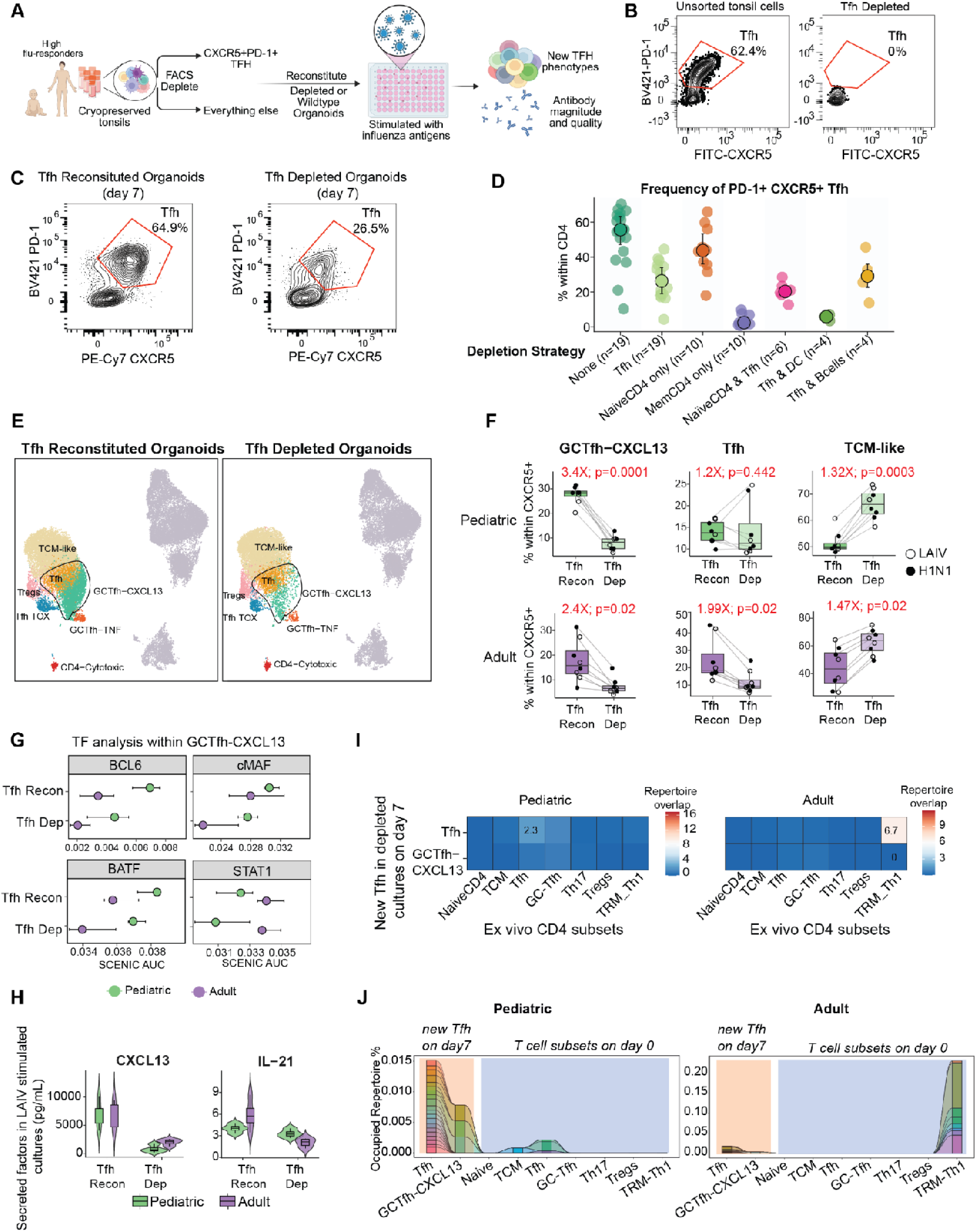
Tfh depletion reveals age-specific clonal origins of newly generated Tfh. (A) Experimental design for Tfh depletion experiments. Tonsil cells were stained and sorted to deplete CXCR5+PD-1+ Tfh. Cells were plated with or without Tfh and stimulated with or without influenza antigens for seven days. Organoids were harvested and B cells and newly generated Tfh were phenotyped using flow cytometry and scRNAseq paired with TCRseq. (B) Representative flow plots demonstrating gating strategy and efficiency of Tfh depletion. (C) Representative flow plots demonstrating the emergence of new Tfh in depleted cultures. (D) Dot plots highlighting frequencies of Tfh in reconstituted and depleted cultures. X-axis denotes the cell type depleted in each experiment relative to reconstituted cultures annotated as “None”. (E) UMAP of and CXCR5+ CD4 T cells and CD19+ B cells from Tfh reconstituted (left) and Tfh depleted organoids (right) from pediatric and adult donors (n=4/group) stimulated with influenza antigens. Clusters of mature Tfh are highlighted. (F) Boxplots comparing proportions of three Tfh subsets within CXCR5+ CD4 compartment in Tfh reconstituted and depleted cultures in pediatric (top) and adult (bottom) donors. Fold change and p-values are highlighted. Boxplots show the median, with hinges indicating the first and third quartiles and whiskers indicating the highest and lowest values within 1.5 the interquartile range. Within each age group, stimulation specific differences were tested using paired Wilcoxon signed-rank tests. (G) Dot plot comparing regulon AUC of targets of key transcription factors in mature GCTfh-CXCL13 subset in flu-stimulated WT and Tfh depleted organoids from pediatric and adult donors. Cell level TF activity scores were aggregated by calculating mean AUC per donor per stim condition. (H) Violin plots comparing secreted levels of CXCL13 and IL-21 in Tfh reconstituted and depleted cultures. Y-axis represents concentrations in pg/mL. Embedded boxplots indicate the median (center line), and interquartile range (box). (I) Heatmap comparing clonal overlap of newly generated Tfh (y-axis) in depleted organoids with ex vivo CD4 T cell subsets (x-axis) in pediatric (left) and adult donors (right) (J) Clonal tracking analysis highlighting top expanded clones in newly generated Tfh in relation to ex vivo CD4 T cell clones from pediatric (left) and adult (right) donors.

To our surprise, new Tfh emerged in Tfh-depleted cultures in significant numbers as early as 24 hours (**fig S5A**), reaching approximately 50% of the frequencies observed in non-depleted cultures (**Figs 6C and 6D**) by day 7 in culture. To identify the cellular determinants of these newly generated Tfh, we systematically depleted other cell types in organoids, either in isolation or in combination with Tfh. These experiments revealed that new Tfh were derived from the memory rather than naive CD4 T cell pool (**Fig. 6D**) and did not require signals or cytokines from B cells, but instead were dependent on dendritic cells (**Fig. 6D**). However, this differentiation process did not rely exclusively on a single signaling pathway, as blocking conventional T helper differentiation signals such as ICOS, IL-6R, or IL-2 individually failed to inhibit new Tfh generation (**sFig 5B**).

We characterized gene expression in these newly generated Tfh compared to Tfh from non-depleted organoids (**Fig. 6A**). Despite their reduced frequencies, new Tfh exhibited transcriptional diversity comparable to Tfh in undepleted cultures (**Fig. 6E**). Although new Tfh failed to reach native GC-Tfh proportions by day 7, a large percentage of them resembled earlier stages of Tfh differentiation (**Figs 6E, 6F and sFig 5C)**. The propensity for Tfh differentiation was age dependent; resting Tfh frequencies reached normal levels in pediatric organoids but remained significantly lower in adults (**Figs. 6E and 6F**), suggesting lesser or slower Tfh differentiation capacity in adult tissues.

We next examined whether newly generated Tfh had similar properties and functional abilities as Tfh from non-depleted organoids. BCL6 protein levels in GC-Tfh were comparable between depleted and reconstituted cultures across both age groups **(sFig 5D)**. Despite this, gene targets of BCL6 were significantly lower in newly generated Tfh (**Fig. 6G**). While ICOS and CXCR5 protein expression remained unchanged, PD-1 and CD200 were reduced in depleted cultures, with a more pronounced decrease in adults **(sFig. 5D)**, suggesting that these cells may be less capable of providing optimal B cell help. Consistent with this observation, secreted levels of both CXCL13 and IL-21 were significantly lower in Tfh depleted organoids (**Fig. 6H**). Interestingly, cMAF target genes, which respond to TGF-β and IL-6 and promote early Tfh commitment (41), were significantly diminished in new Tfh in adults but not in pediatric donors (**Fig. 6G**), consistent with age-dependent differences in Tfh differentiation pathways.

To investigate age-associated differences in cellular origins of newly arising Tfh, we assessed clonal sharing between new Tfh on day 7 with both Tfh and non-Tfh CD4 clones from the same donors *ex vivo* (**Figs. 6I and 6J**). This analysis revealed striking age-dependent differences in Tfh differentiation potential. In pediatric donors, new Tfh arose predominantly from central memory CD4 T cells (TCM) that were clonally related to ex vivo Tfh. In contrast, new Tfh in adults were clonally related to effector-like CD4 T cells, suggesting distinct differentiation pathways between age groups (**Figs 6I and 6J**). In conclusion, these findings provide a mechanistic basis for age-dependent differences in Tfh differentiation and maturation profiles in the lymphoid tissue microenvironment.

### Age-dependent effects of Tfh depletion on B cell responses

Finally, we examined how Tfh depletion altered GC selection, and antibody production and quality in pediatric vs. adult donors. To our surprise, the magnitude of influenza HA-specific antibodies was unchanged by Tfh depletion in both pediatric and adult cultures (**Fig. 7A**). However, virus microneutralization assays revealed a marked deficit in antibody quality exclusively in adults (**Fig. 7B**), suggesting that pediatric donors might rely less on canonical Tfh help to generate functional influenza antibodies.

**Fig 7:**
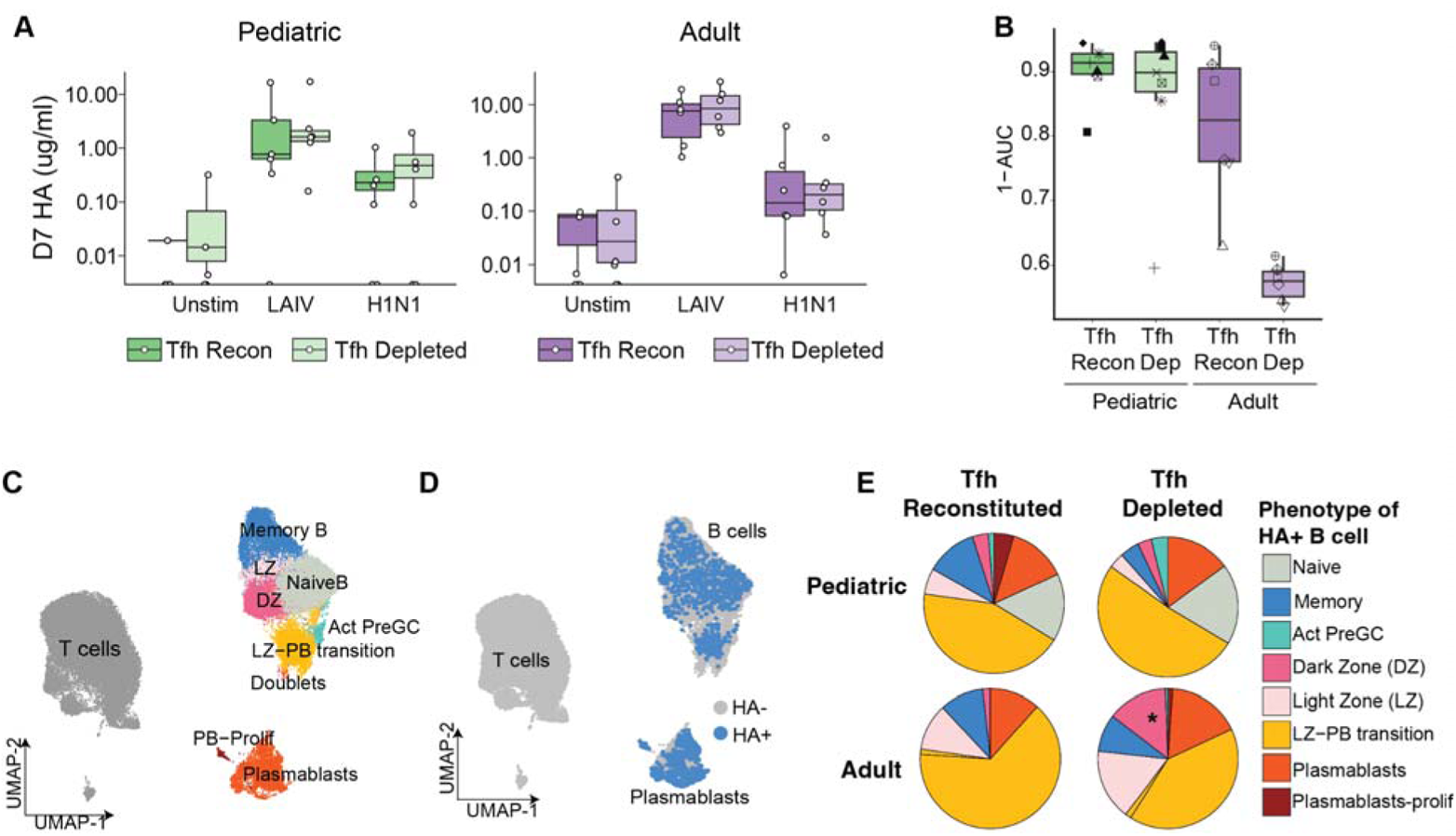
Age-dependent differences in the requirement for Tfh support during GC maturation and antibody production. (A) Boxplots comparing influenza specific antibodies (HA-specific, in ug/mL) in Tfh reconstituted and depleted organoids generated from pediatric (left) and adult (right) donors. (B) Comparison of virus microneutralization data, quantified by area under the curve (AUC) for Tfh reconstituted and depleted organoids stimulated with LAIV in pediatric and adult donors. Boxplots show the median, with hinges indicating the first and third quartiles and whiskers indicating the highest and lowest values within 1.5 the interquartile range. (C-D) UMAP of and CXCR5+ CD4 T cells and CD19+ B cells from wildtype (left) and Tfh depleted organoids (right) from pediatric and adult donors (n=4/group) stimulated with influenza antigens. Clusters of all B cells are highlighted in (C), and antigen (HA) specific B cells are overlaid in (D) (E) Pie charts illustrating distribution of HA+ B cells in Tfh reconstituted (left) and depleted (right) organoids stimulated with LAIV in pediatric (top) vs. adult (bottom) donors. HA cells within dark zone (DZ) are highlighted with *.

To define the cellular basis for this defect, we quantified and phenotyped HA+ B cells in Tfh depleted and reconstituted organoids using flow cytometry and scRNAseq **(Figs 6A and sFig 6A)** as previously described (28,39). Overall frequencies of HA+ cells with a GC or plasmablast phenotype did not change with Tfh depletion **(sFig 6B)**, but their transcriptomic profiles (**Figs. 7B and 7C**) revealed differences in cell state. In adult organoids, Tfh depletion resulted in accumulation of HA+ B with a Dark Zone GC phenotype (**Fig 7E and sFig 6C)**. Together, these data show that early life influenza antibody responses can be maintained in a Tfh independent manner, whereas adult B cell responses to influenza antigens are reliant on selection cues from Tfh for affinity maturation. This Tfh-dependent failure of selection likely underlies the sharp decline in neutralizing capacity observed in adult cultures.

## DISCUSSION

In this study, we leveraged mucosal tissue sampling from *in vivo* vaccinated and not recently vaccinated donors, human tonsil organoids, functional assays, and single cell genomics to define the phenotypic, functional, and differentiation properties of Tfh across early life and adulthood. Overall, our findings reveal that while pediatric tonsils harbor abundant, diverse Tfh subsets with high clonal diversity and plasticity, they exhibit fundamental differences in functional maturity, activation programs, and reliance on canonical GC-associated signaling compared to their adult counterparts. Critically, we demonstrate that pediatric Tfh are not merely immature versions of adult Tfh but rather operate within a distinct immune framework optimized for the unique immunological challenges of early life (22,42).

A central finding of our study is that diverse Tfh subsets are more abundant in pediatric tonsils than in adults (26), yet these cells express lower surface expression of critical B cell help molecules (CD40L, OX40, CD84) and produce substantially less IFNL, TNFα, and IL-17 following TCR stimulation. This disconnect between Tfh abundance and functional output challenges previous assumptions that Tfh frequency predicts the quality of humoral responses. Interestingly, we observed higher transcriptionally defined Tfh-IL10 and Tfh-TNF subsets in young children at baseline, but these cytokines were secreted at lower levels following stimulation compared to adult Tfh. Adult tonsils also harbored more Tfh1 and Tfh17 cells compared to young children, consistent with previous observations of circulating Tfh (cTfh) in blood (7). This apparent paradox suggests that early-life Tfh are in a poised but incompletely activated state. This most likely represents an adaptive mechanism to prevent excessive inflammation in tissues continuously exposed to commensal and environmental antigens, though at the cost of reduced helper capacity during vaccine responses (43,44). The enhanced clonal diversity and plasticity we observed in early-life Tfh further support the notion of a less differentiated, more flexible state. Cytokine-expressing Tfh subsets exhibited substantial clonal overlap with GC-Tfh in children but not adults, suggesting that individual clones can more readily transition between phenotypic states. This plasticity may be advantageous for building a diverse immune repertoire during early antigen encounters (45) but could compromise the stable, polarized effector programs required for optimal GC B cell selection.

Beyond quantitative differences in cytokine production, we identified qualitatively distinct transcriptional programs activated by pediatric versus adult Tfh after influenza immunization. Specifically, GC-Tfh responding to vaccination mounted strong type-I interferon responses in pediatric donors, suggesting a heightened antiviral state (46). In contrast, adult GC-Tfh exhibited robust antigen-specific effector programs, upregulated Th1 associated transcription programs and polarized cytokine production. Specifically, following influenza vaccination, BCL6 targets in GC-Tfh were downregulated while Blimp-1 targets were upregulated prominently in adults, indicating progression toward an effector state. This regulatory switch was absent in pediatric GC-Tfh, suggesting they maintain a less differentiated state that may account for their sustained yet weak effector responses. This strategy may be adaptive in early life, when innate-like responses provide rapid and broad protection against a barrage of new pathogens (22), even if these responses come at the cost of the precision and durability characteristic of adult immunity.

A critical unresolved question we sought to address in this study is whether poor pediatric vaccine responses reflect intrinsic defects in Tfh, or simply insufficient antigen experience. Our previous immune organoid studies have shown that a subset of pediatric donors mount robust neutralizing antibody responses to LAIV, comparable to those in adults (30). Restricting our analysis to these high-responding donors, we observed that pediatric Tfh achieved antigen-specific activation levels and clonal expansion comparable to those of adults, with activated Tfh1 frequencies correlating with neutralizing antibody quality regardless of age. These findings demonstrate that pediatric Tfh possess the intrinsic capacity for antigen-specific activation and proliferation in high antibody-producing donors and suggests that optimizing responses to activate Tfh1 cells, rather than simply increasing Tfh numbers, might be a good strategy to enhance vaccine responses in young children.

Recent work in murine models have identified CD62L+ stem-like memory cells that efficiently generate Tfh with effector functions in recall responses (47). Our Tfh depletion experiments in organoids revealed age-dependent differences in both the differentiation capacity and the origins of newly generated Tfh. While new Tfh failed to acquire fully functional GC-Tfh phenotypes in both groups, the resting Tfh population returned to baseline levels and was populated by differentiation of central memory CD4 T cells. In contrast, adult Tfh emerged from effector-like CD4 T cells, indicating recruitment from a more differentiated memory pool. The ability of pediatric central memory cells to efficiently regenerate Tfh may represent an adaptive mechanism to rapidly build or rebuild GC responses following repeated mucosal exposures. Surprisingly, blockade of individual signaling pathways previously shown to be critical for Tfh differentiation (ICOS, IL6R, or IL-2) failed to prevent new Tfh generation. This indicates that multiple mechanisms can drive Tfh differentiation and that the process is robust to depletion of a single signaling factor, likely reflecting redundancy in the priming signals provided by dendritic cells and other antigen-presenting cells (48) in organoid cultures. However, this redundancy raises questions about what pathways to target therapeutically to enhance pediatric Tfh function.

The most striking finding of our study is that pediatric B cell responses to influenza vaccination are substantially less dependent on Tfh compared to adults. This functional divergence was mirrored at the cellular level: Tfh depletion retained HA-specific cells in the GC dark zone but did not affect pediatric GCs. Moreover, IL-21 secretion was significantly reduced in Tfh-depleted adult organoids, indicating that pediatric GC B cells function independently of IL-21 or utilize alternative survival signals, bypassing Tfh-dependent selection. Reduced IL-21 reliance appears to be a conserved feature of early life immunity, as murine models of hepatitis B infection similarly show diminished IL-21 during early life, which contributed to impaired adaptive immunity and viral persistence (49). Consistent with this, cell-cell interaction analysis revealed weaker Tfh-B cell interactions at key GC checkpoints (IL21R, CD80, TACI) in LAIV-stimulated organoids. Furthermore, ICOS blockade, which attenuated both Tfh1 and IL-21 responses in adults, had minimal impact on pediatric Tfh function. Together, these findings suggest that while adults require canonical ICOS and IL-21 mediated Tfh help for high-affinity antibodies, pediatric GCs might employ less stringent selection thresholds, allowing moderate-affinity B cells to differentiate into plasmablasts via extrafollicular or other alternative pathways (50).

Our findings have broad implications for pediatric vaccine design. First, they suggest that simply increasing Tfh frequency or broad activation, for example through adjuvants targeting Tfh differentiation, will be insufficient if those Tfh lack functional maturity. Strategies that promote Tfh polarization towards Th1-like phenotypes and enhance IL-21 production may be more effective for enhancing humoral responses. Adjuvants that engage TLRs and drive type-I interferon responses are already used in some pediatric vaccines, but our data suggest they may promote innate-like rather than antigen-specific Tfh programs. Alternative adjuvants that drive STAT1/STAT4 signaling and support TBET expression warrant investigation.

Our data also show that pediatric Tfh with adequate pre-existing memory are functionally competent compared to adults. This supports the use of multi-dose regimens for building pediatric immunity through vaccination. However, if pediatric B cells are adapted to rapidly produce antibodies in a Tfh independent manner, interventions that force stringent Tfh-dependent selection might delay protective immunity without commensurate gains in affinity. Future longitudinal studies tracking antibody durability and recall responses will be essential to determine whether Tfh-independent pediatric responses provide adequate long-term protection. Some limitations of our study warrant consideration. First, we focused on tonsils as a tractable GC-rich tissue, but whether our findings generalize to other lymphoid sites, particularly the draining lymph nodes close to the sites of vaccination, is yet to be determined. Second, our pediatric cohort consisted of relatively healthy 2-3 year olds undergoing elective tonsillectomy. Tfh biology in neonates, where vaccine failures are most severe, may differ substantially. Third, organoid cultures lack the full systemic context of *in vivo* immunity, including circulating antibodies, complement, and immune cells recruited from distal lymphoid tissues. While there was some degree of agreement between *in vivo* and *in vitro* Tfh responses to influenza vaccination, more extensive clinical studies would be needed to validate our observations. Nevertheless, understanding age-dependent differences in Tfh biology is essential for designing vaccines that work with, rather than against, the unique features of early-life immunity. Ultimately, our findings position Tfh as a promising target for improving pediatric immune responses and offer a rational framework for closing the immunity gap that leaves young children vulnerable to preventable infections.

## MATERIALS AND METHODS

### Informed consent and sample collection

Tonsils from healthy consented individuals undergoing surgery for obstructive sleep apnea, hypertrophy, or recurrent tonsillitis were collected in accordance with the University of California, Irvine Institutional Review Board (IRB). Ethics approval was granted by the University of California, Irvine IRB (protocol #2020-6075), and all participants provided written informed consent. Participants in the primary cohort used for the study were aged 2-58 years (Table S1). Overall, tonsil tissue was healthy in appearance, and we have not observed differences in cell composition or organoid responses related to surgical indication.

Recently vaccinated donors were enrolled in the study after recruitment from the ENT outpatient clinic at Haukeland University Hospital, Norway. All subjects were patients scheduled for elective tonsillectomies (tonsillitis, hypertrophy, or both) but otherwise healthy. Written informed consent was obtained from all donors (NCT01866540). Donors were vaccinated with trivalent LAIV (Fluenz; AstraZeneca) 5-9 days before tonsillectomies.

### Sample processing

Samples were processed as previously described (39). Briefly, whole tonsils were collected in saline after surgery and immersed in an antimicrobial bath of Ham’s F12 medium (Gibco) containing Normocin (InvivoGen), penicillin, and streptomycin for 30-60 min at 4°C for decontamination of the tissue. Tonsils were then briefly rinsed with PBS and mechanically dissociated; debris was removed using gradient centrifugation (Lymphoprep, Stemcell). Cells were counted, and samples were cryopreserved in fetal bovine serum (FBS) with 10% DMSO and stored in nitrogen until use.

### Immune Organoids

To generate organoids, cryopreserved cells were thawed, enumerated, and then plated at a final density of 7.5 x10^6^/ml (200 μl final volume in ultra-low attachment plates (Corning)). Organoid media was composed of RPMI1640 with glutamax, 10% FBS, 1x nonessential amino acids, 1x sodium pyruvate, 1x Antibiotic-Antimycotic (Gibco), 1x Normocin (InvivoGen), 1x insulin, selenium, transferrin supplement (Gibco), and 0.5 μg/ml recombinant human BAFF (BioLegend). Antigen doses were previously titrated (28) and dosing was selected based on the ability to induce B cell activation and antibody production without affecting organoid viability. The following amounts of influenza antigens were added to immune organoids at culture - 1:2,000 final dilution of 2019/20 live attenuated influenza vaccine (LAIV, FluMist® Quadrivalent) and 2.5 hemagglutination units (HAU) per culture of A/California/07/2009 H1N1 virus. Cultures were incubated at 37°C, 5% CO2 with humidity for two weeks (unless otherwise stated in the experimental design) and media was replenished every other day by exchanging 30% of the volume with fresh organoid media.

### Cell Sorting

Thawed tonsil cells were stained, and bulk sorted into culture medium containing 20% FBS using a FACSAria Fusion. Sorting experiments involved separating individual cell subsets or combinations in some case, with the following markers: Tfh (CXCR5++ PD-1++); Naive CD4 (CD4+ CD45RA+); Memory CD4 (CD4+ CD45RA-); B cells (CD3- CD19+); DCs (CD3- CD19-HLA-DR+ CD11c+ CD36+). Cell types were depleted by FACS from day 0 tonsil cells and cultured with LAIV or H1N1 for 7 days as described in the “Immune Organoids” section. As controls, depleted cultures were reconstituted with the cell type that was originally sorted and plated at the same cell density as the depleted cultures. A post-sort analysis was used to ensure depletion purity.

### Flow Cytometry

Tonsillar cells or harvested immune organoids were washed with FACS buffer (PBS + 0.1% BSA, 0.05% Sodium Azide, and 2 mM EDTA) to remove any residual antibodies or factors generated during culture. All samples were stained with antibody cocktails (Table S2) in FACS buffer for 30 minutes on ice while protected from light. Data were collected using a Quanteon ACEA Novocyte or Cytek 5L Aurora flow cytometer.

### Antigen specific B cell staining

A/California/07/2009 (H1N1) hemagglutinin protein (Immune Technology) was biotinylated per manufacturer’s instructions using the EZ-link micro-NHS-PEG4-biotinylation kit (Thermo Scientific) and as described previously (51). Protein was incubated with 50 mmol excess biotin at room temperature (RT) for 45 minutes and buffer was exchanged using zeba spin desalting column to remove excess biotin. For staining of HA+ B cells, organoids were harvested as described above and first incubated with biotinylated HA (2μg/ml final concentration) and Human TruStain FcX (Biolegend) for 30 minutes on ice. Cells were thoroughly washed with FACS buffer and stained with a cocktail of cell surface antibodies and streptavidin-PE and streptavidin-APC conjugates to detect antigen-specific cells. Cells were washed with FACS buffer and analyzed on Quanteon ACEA Novocyte or sorted on FACSAria Fusion.

### Intracellular cytokine staining

Immune organoids were harvested and washed with FACS buffer. Samples were surface stained with antibody cocktails prepared in FACS buffer for 30 minutes on ice while protected from light. Cells were washed with FACS buffer and fixed with 1X Cyto-Fast Fix/Perm buffer (BioLegend) at RT for 30 minutes in the dark. Pellets were washed twice with 1X Cyto-Fast Perm Wash buffer and stained with a cocktail of intracellular antibodies prepared in 1X Cyto-Fast Perm Wash buffer for an hour at RT in the dark (Table S2). Cells were then washed once with 1X Cyto-Fast Perm Wash buffer and twice with FACS buffer, resuspended in FACS buffer, and analyzed on Cytek 5L Aurora flow cytometer. Fluorescence Minus One (FMO) controls for each cytokine served as controls for gating of cytokine+ cells.

### Intranuclear staining

For intranuclear staining of Bcl6 and Blimp1, ex vivo tonsil cells or harvested organoids were surface stained as described above. Cells were then washed with FACS buffer and fixed and permeabilized for intranuclear staining for 1 h at 25C with True Nuclear Fixation buffer (BioLegend), washed twice with permeabilization/wash buffer, and stained for 1 hr at 25C with Bcl-PE and Blimp1-APC. Cells were washed twice with permeabilization/wash buffer and resuspend in FACS buffer for acquisition on Cytek 5L Aurora flow cytometer. Fluorescence Minus One (FMO) controls for each transcription factor served as controls for gating of positive cells.

### Antibody detection by ELISA

Influenza-specific antibodies were detected as previously described. High-binding assay plates (Corning) were coated overnight with influenza proteins at a final concentration of 2 ug/mL in 100 mM sodium bicarbonate coating buffer. Non-specific binding was blocked by incubating plates with 1% BSA in PBS for 2 hours at RT. Culture supernatants were diluted 1:20 (for unstim) or 1:80 (for LAIV or H1N1 stimulated cultures) in PBS and added to coated, blocked plates for 1 hour at RT. Horseradish peroxidase-conjugated anti-human secondary antibodies to IgM/IgG/IgA (Abcam) were used to detect bound antibodies. A monoclonal influenza IgG antibody (clone CR9114) was used as a standard to estimate HA-specific antibody concentrations.

### Microneutralization assays

Microneutralization assays (MN) in this study were conducted as previously described. Briefly, Madin-Darby canine kidney cells (MDCKs, ATCC) were cultivated in Eagle’s minimum essential medium (EMEM, ATCC) supplemented with 1% Anti-Anti and 10% heat-inactivated FBS and incubated at 37°C with 5% CO2. One day prior to the assay, MDCK cells were subcultured into flat-bottomed 96-well plates at a density of 1.1 × 10^4^ cells in 100μL per well. Culture supernatants from Tfh depleted and reconstituted organoids were diluted (1/2.5) in virus growth media (VGM) consisting of serum-free EMEM supplemented with 0.6% bovine serum albumin (BSA, Sigma-Aldrich) and 1 μg/mL N-p-Tosyl-l-phenylalanine chloromethyl ketone (TPCK)-treated trypsin (Worthington Biochemical) and then serially diluted (two-fold) in VGM, 50μl were left over in all wells. The infectious A/California/07/09 X A/Puerto Rico/8/1934 reassortant H1N1 virus (BEI NR-44004) was diluted to 50 TCID50 per 50 μL in VGM and subsequently added to the serially diluted supernatants, followed by an incubation period of 1 hour at 37°C and 5% CO2. Replicate control samples consisting of 100μl diluted virus only or 100μl VGM only were also prepared. After incubation, the media from MDCK monolayers was replaced with the serum-virus mixtures and further incubated for 1 hour. Following this, the serum-virus mixtures were substituted with 200 μL of VGM supplemented with 2% FBS, and cells were incubated for 48 hours at 37°C. Post-incubation, media was removed, and cells were fixed in 4% paraformaldehyde (PFA) in PBS for 30 minutes, washed once in PBS, and then permeabilized in 0.1% PBS/Triton X-100 (PBS-T) at room temperature for 15 minutes. Next, cells were washed twice with PBS and blocked using 200μl of blocking buffer (3% BSA) in PBS for 1 hour at room temperature. Influenza virus nucleoprotein (NP) was detected utilizing an equal mixture of anti-NP mAbs (Millipore Cat Nos. MAB 8257 and MAB 8258) diluted (1/1000) in blocking buffer, followed by horseradish peroxidase (HRP)-conjugated anti-mouse IgG (KPL, Cat. No. 074-1802) diluted at 1:3000 in blocking buffer. Plates were developed in TMB peroxidase substrate and reactions were halted using 1N HCl. Finally, assays were quantified in an ELISA plate reader at 450 nm using SoftMax Pro 7.1 software.

### Luminex

Secreted cytokines, chemokines, and growth factors from day 6 organoids culture supernatants were measured using a custom 36-plex human cytokine Luminex assay combined with a 3-plex kit (Millipore Sigma). Frozen supernatants that were previously never thawed were diluted 1:2 and plated in duplicates. Plates were processed per manufacturer’s instructions, modified to analyze 384 samples (instead of 96) on the MAGPIX or Intelliflex multiplexing system. Standard curves were generated using five-parameter logistic regression on the Belysa (Millipore Sigma) or xPONENT software (Luminex Corp).

### scRNA-seq of ex vivo tonsil CD4 T cells

Cryopreserved tonsils from donors with or without recent influenza vaccination were revived using pre-warmed RPMI1640 media supplemented with glutamax, 10% heat-inactivated FBS, 1x nonessential amino acids, 1x sodium pyruvate, and 1x penicillin-streptomycin. Cells were washed with FACS buffer, and roughly 1-2 10^6^ cells were stained with a cocktail containing fluorescent labeled antibodies for sorting made in FACS buffer for 30 minutes on ice in the dark. Each donor/sample received a unique hashing antibody (TotalSeqC, BioLegend) for sample barcoding. Following the 30 min incubation, cell pellets were resuspended in 200 uL FACS buffer and placed on ice until sort. Roughly 50K T cells from each donor were sorted into the same collection tube based on viable single cells with markers CD19-CD3+. Pooled cells were then pelleted and stained with a cocktail of DNA oligo-tagged cell surface antibodies (Table S2) for 30 minutes on ice. Cell pellets were then washed thoroughly in 4 mL FACS buffer for a total of 4 washes. After the final wash, pellets were resuspended in 100 μl ice-cold wash buffer (PBS + 0.04% BSA), counted, and resuspended to a final concentration of 2,400 cells/uL. Single-cell suspensions were superloaded on a 10X Genomics Chromium controller with a loading target of 60,000 cells per sample. Libraries were generated using the Chromium Next Gem Single Cell 5’ Reagent Kit v2 (Dual Index) per the manufacturer’s instructions with the addition of 5’ Feature Barcode libraries. Quality and quantity of libraries were measured on tapestation, qubit, and bioanalyzer, and sequenced on Illumina NovaSeq 6000 with a sequencing target of 30,000 reads per cell for gene expression libraries and 10,000 reads per cell for TCR and feature barcode libraries.

### scRNA-seq of immune organoids

Tonsil immune organoids from both sorted and unsorted (Tfh depletion) experiments were harvested, washed in FACS buffer and stained with biotinylated HA protein as described above. Cells were washed three times, and surface stained with a cocktail containing fluorescently labeled antibodies for sorting, strepatvidin, in addition to DNA oligo-tagged cell surface antibodies (TotalSeq-C, Table S2). Additionally, each organoid was labeled with a unique hashing antibody (hashtag oligos, HTO), allowing for high throughput sample multiplexing. Samples were stained for 30 minutes in the dark on ice, washed three times with FACS buffer, and left on ice until sorting. For immune organoids in Fig 3, samples were washed with FACS buffer and sorted to enrich total CD4 T cells and HA+ B cells into one collection tube, and HA- B cells into another. For immune organoids from Tfh depleted cultures in Figs 6 and 7, samples were washed with FACS buffer and sorted to enrich CXCR5+ CD4 T cells and HA+ B cells into one collection tube, and HA- B cells into another. In both experiments, sample hashing with HTO allowed us to pool several organoids into one tube. Pooled samples were pelleted, and cells were resuspended in 100 μl ice-cold wash buffer (PBS + 0.04% BSA), counted, and resuspended to a final concentration of 2,400 cells/uL. Single-cell suspensions were then immediately loaded on a 10X Chromium Controller with a loading target of 60,000 cells. Libraries were generated using the Chromium Next Gem Single Cell 5’ Reagent Kit v2 (Dual Index) per the manufacturer’s instructions with the addition of 5’ Feature Barcode libraries. Quality and quantity of libraries were measured on tapestation, qubit, and bioanalyzer, and sequenced on Illumina NovaSeq 6000 with a sequencing target of 30,000 reads per cell for gene expression libraries and 10,000 reads per cell for TCR and feature barcode libraries.

### 5’ gene expression analysis

#### Data Alignment

Raw reads from gene expression, TCR, and HTO libraries were aligned and quantified using Cell Ranger Single Cell Software Suite with Feature Barcode addition (10X Genomics) against GRCh38 human reference genome. Alignment was performed using feature and vdj option (vdj_GRCh38_alts_ensembl_5.0.0) in cellranger. Feature files from Cell Ranger were manually updated to separate hashing antibodies (HTO) from other cell surface antibodies (Protein) to facilitate independent normalization procedure.

#### TCR gene suppression

For each sample, the gene expression matrix was loaded in Seurat (version 5.0.2) and TCR genes were summarized into custom *TCRA*, *TCRB*, *TCRG*, and *TCRD* genes to avoid downstream clustering based on V/D/J gene usage. Cellular features such as mitochondrial and ribosomal gene expression and the number of housekeeping genes were added as additional meta-features.

#### Doublet removal and QC

Within each library, doublets were identified computationally, with an expected doublet rate of 0.4% per 1,000 cells. DoubletFinder (version 3) was run on default settings (pN = 0.25, pK = 0.09, PCs=1:20) and removed from subsequent QC steps. Poor quality cells and droplets with ambient RNA (number of expressed genes less than 200 and housekeeping genes less than 25) and additional doublets (number of expressed genes more than 4000) were excluded from subsequent analysis.

#### Hash Demultiplexing

Sample assignment and further doublet removal was performed using HTODemux function using default settings on Seurat’s HTO vignette.

#### Normalization and Clustering

Samples from each experiment were merged using Seurat’s *merge* function. In Fig 3, pediatric organoids were integrated with previously published organoid data from adults, with the exclusion of a stimulation condition (Inactivated Influenza Vaccine). Data normalization and variance stablization were performed on the merged object using NormalizeData and ScaleData functions in Seurat. Dimensionality reduction was performed using *RunPCA* function to obtain the first 30 principal components, of which the first 20 were used for clustering using Seurat’s *RunUMAP* function with a 0.8 resolution. Cell types were broadly assigned as *CD3D* expressing T cells or *CD19* or *MS4A1* expressing B cells, which were further subsetted for reclustering as described above. Contaminating B cell clusters from subsetted T cell clusters were iteratively removed and renormalized until the final UMAP contained only CD4 T cells.

#### Batch Correction

Batch variability between libraries generated in batches were corrected using Harmony using default settings in the vignette.

#### Cluster Annotation

Gene and protein markers for each cluster were identified using *FindMarkers* function in Seurat. For gene expression, we used Wilcoxon Rank Sum tests for differential marker detection using a log2 fold change cutoff of 0.4. CD4 T cell clusters were identified using a known catalog of markers defined for T cells from human tonsils (23,36). For cell surface protein, data were normalized using centered log-ratio (CLR) transformation. A combination of both positive and negative protein markers from each cluster was used to finalize annotations. A final list of markers is provided in Table S3.

#### Pseudobulk and Pathway analysis

Pseudobulk gene expression profiles was generated using Seurat’s *AggregateExpression* function, aggregating raw RNA counts by donor, age group, and Tfh cluster. Differential expression for pseudobulk samples was analyzed using DESeq2. Genes were ranked by log2 fold change for subsequent pathway analysis using gene sets from Molecular Signatures Database (MSigDB) accessed through msigdbr. Ranked gene lists were tested against seven pathway collections: Hallmark gene sets (H), BioCarta (C2:CP:BIOCARTA), KEGG (C2:CP:KEGG), PID (C2:CP:PID), Reactome (C2:CP:REACTOME), WikiPathways (C2:CP:WIKIPATHWAYS), and Gene Ontology Biological Process (C5:GO:BP). Enrichment analysis used gene set size filters of 15-500 genes and 1,000 permutations. Pathways were ranked by normalized enrichment score (NES), and statistical significance was assessed at adjusted p-value < 0.05.

#### Transcription factor analysis

To identify active transcription factors (TFs) and their target regulons, we performed single-cell regulatory network inference and clustering (SCENIC) using the pySCENIC pipeline (52). Co-expression modules between TFs and potential target genes were inferred using GRNBoost2, followed by cis-regulatory motif enrichment analysis to confirm direct regulatory interactions using the hg38 motif database (10kbp_up_10kbp_down and 500bp_up_100bp_down) and HGNC transcription factor list. Regulons were defined as TFs with a Normalized Enriched Score of at least 1.5. Activity of these regulons within each cell was quantified by the AUCell algorithm. To assess differential TF activity between age groups for key Tfh associated regulators, cell-level regulon activity scores were aggregated by calculating mean AUC per donor and testing significance using unpaired t-tests with Bonferroni correction.

#### Ligand Receptor analysis

To infer cell-cell communication networks using transcriptomic data from vaccine stimulated immune organoids, we utilized the MultiNicheNet framework (53). Analysis was restricted to sender and receiver clusters present in both age groups under unstimulated and LAIV conditions, with samples aggregated by donor. Multi-level differential expression analysis was performed to compare LAIV treatment relative to unstim controls in each age group independently. Pseudobulk differential analysis was performed for ligands in sender cells and receptor/target genes in receiver cells (minimum 10 cells, fraction cutoff 0.05, p < 0.05, log2FC > 0.5). Ligand activity was predicted by assessing enrichment of downstream target genes using the NicheNet ligand-target matrix. Prioritized ligand-receptor pairs were ranked using MultiNicheNet’s “regular” scoring scenario integrating ligand activity, expression, and differential expression scores with top 50 interactions extracted per contrast.

### TCR analysis

Only QC-filtered cells with confirmed annotations were included in downstream TCR analysis. Basic features of the repertoire (clonality by donor, by cluster, stimulation condition) was described using standard workflows in Immunarch (54). Clonal overlap and tracking analysis for comparison of ex vivo clusters with T cell clusters from immune organoids was performed using *repOverlap* (morisita overlap) and *trackClonotypes* function in immunarch.

### Statistical analysis

All statistical analyses were performed in R. For comparing paired samples across stimulation conditions or time points, we used paired Wilcoxon signed-rank tests (two-sided). For comparing age-differences, unpaired Mann-Whitney U tests (two-sided) were performed.

## RESOURCE AVAILABILITY

Requests for further information or access to data should be directed to Lisa Wagar

## MATERIALS AVAILABILITY

This study did not generate new unique reagents.

## DATA AVAILABILITY

Raw data has been deposited on NCBI’s Sequence Read Archive (project ID: pending)

## ACKNOWLEDGEMENTS

The authors thank the tonsillectomy patients and their families for participating in this study, Drs. Jennifer Atwood, Michael Hou, Pauline Nguyen, and Vanessa Scarfone from UC Irvine flow cytometry cores for technical assistance with flow sorting, Drs. Robert Edwards and Delia Tifrea at the University of California, Irvine Medical Center pathology for sample coordination, and Dr. Melanie Oakes at UC Irvine Genomics Research and Technology Hub (GRT-Hub) for assistance with library preparation.

This work is supported by funding from the Wellcome Leap HOPE Program (to L.E.W) and the National Institutes of Health (NIH) grant R01AI173023 (to L.E.W). S.S. is supported by the National Institutes of Health K99/R00 Pathway to Independence Award (K99AG090739) funded by the National Institute on Aging (NIA). This work was also made possible, in part by the Genome Technology Access Center at the McDonnell Genome Institute at Washington University School of Medicine. The center is partially supported by NCI Cancer Center Support Grant P30 CA91842 to the Siteman Cancer Center from the National Center for Research Resources (NCRR), a component of the National Institutes of Health (NIH), and NIH Roadmap for Medical Research. This work is supported by funding from the Children’s Discovery Institute of Washington University and St. Louis Children’s Hospital (MI-LI-2020-914 to N.S.).

This work was also made possible, in part, through access to the following: the Genomics Research and Technology Hub (formerly Genomics High-Throughput Facility) Shared Resource of the Cancer Center Support Grant (P30CA-062203), the Single Cell Analysis Core shared resource of Complexity, Cooperation, and Community in Cancer (U54CA217378), the Genomics-Bioinformatics Core of the Skin Biology Resource Based Center (P30AR075047) at the University of California, Irvine and NIH shared instrumentation grants 1S10RR025496-01, 1S10OD010794-01, and 1S10OD021718-01. This publication is solely the responsibility of the authors and does not necessarily represent the official view of the funding agencies.

## AUTHOR CONTRIBUTIONS

Designed, optimized, and performed experiments: SS, JMK, EMJ, ZWW, SK, NZ, GB, AM, GM, CT, MTM, AD, CT, FZ, KAB

Analyzed and interpreted data: SS, JMK, EMJ, AKN, SK, NZ, TBY, EC

Sample procurement: AMS, GM, HB, DT, GA, QZ, RJC

Conceived the study, study design, and secured funding: SS, JMK, NS, RJC, LEW Writing the original manuscript: SS and LEW

Edited, reviewed, and approved manuscript: all authors

## DECLARATION OF INTERESTS

JMK is currently an employee of F. Hoffman La Roche, Basel, Switzerland. LEW declares inventor status on a US patent (US-20230235284-A1) describing the immune organoid technology. The other authors declare no competing interests.

### Supplementary Materials

sFig 1: scRNA-seq profiles of tonsillar CD4 T cell subsets in pediatric and adult donors

sFig 2: Comparing Tfh phenotypes from pediatric donors and adults

sFig 3: scRNA-seq profiles of flu-stimulated tonsil organoids

sFig 4: Correlating Tfh responses in organoids with antibody responses

sFig 5: Tfh depletion strategy in tonsil organoids

sFig 6: Characterization of B cell responses following Tfh depletion

Table S1: Cohort of human donors used in this study

Table S2: List of feature barcode antibodies used in this study

Table S3: List of cluster-specific top gene expression markers for all single cell experiments in the study.

