## Supplementary material for "Divergent phenotypic and functional roles of human T follicular helper cells from infancy to adulthood": Supp Figures

fig S1

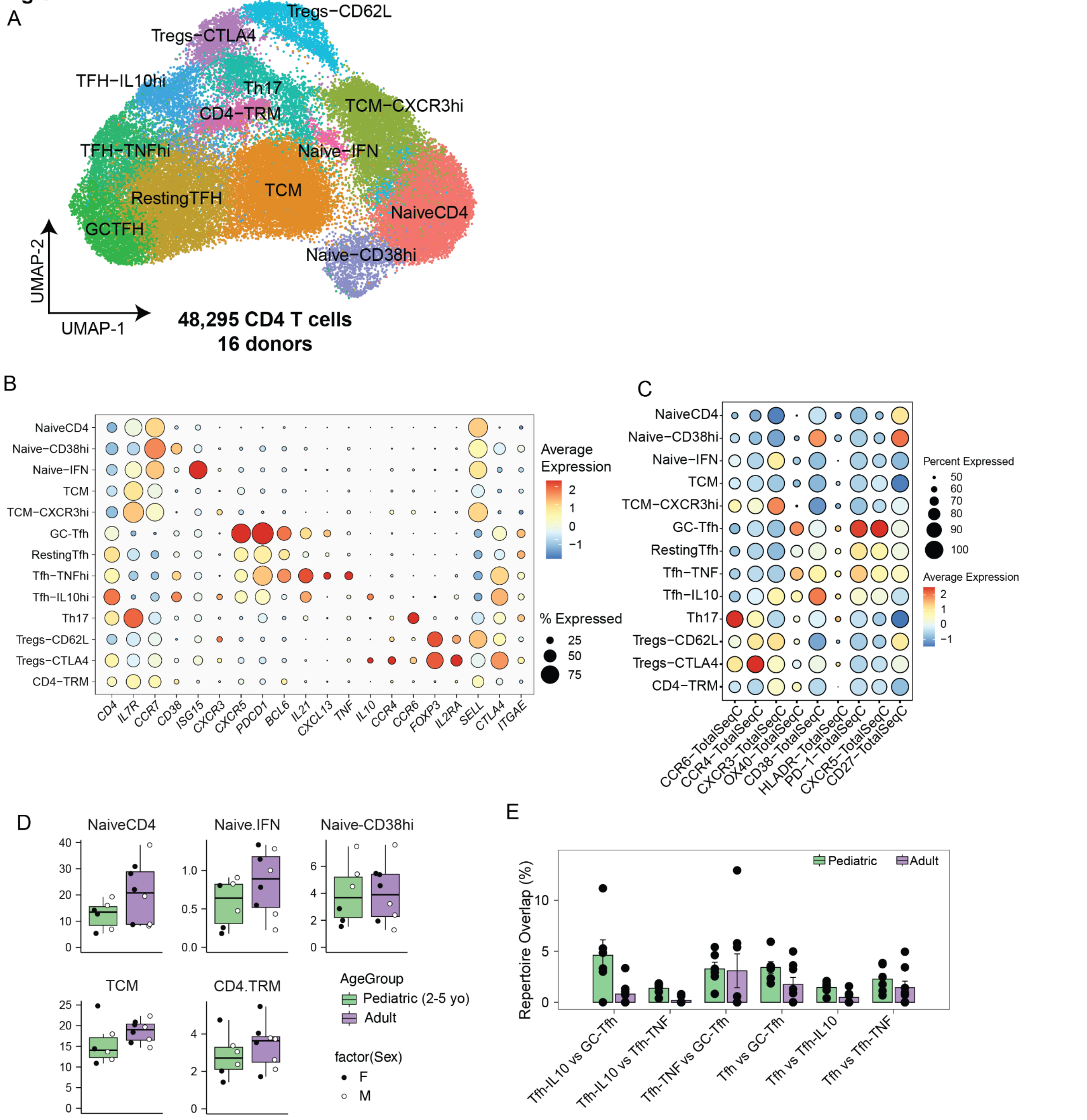


**sFig 1: scRNA-seq profiles of tonsillar CD4 T cell subsets in pediatric and adult donors.** (A) UMAP of 48295 CD4 T cells from palatine tonsils from 12 donors (n=6/group) (B) Bubble plot comparing gene expression markers among CD4 subsets in the tonsils. (C) Bubble plot comparing surface protein markers among CD4 subsets in the tonsils. The size of the bubble indicates the proportion of cells within the cluster expressing the marker, and the color indicates the magnitude of expression ranging from blue (low) to red (high). (D) Boxplots comparing proportions of non Tfh subsets within the total CD4 compartment in tonsils from pediatric (<5 years old, n=6) and adult donors (n=6). (E) Dotplots comparing clonal overlap among Tfh subsets in pediatric and adult donors. Clonal overlap was calculated using immunarch.

**fig S2**

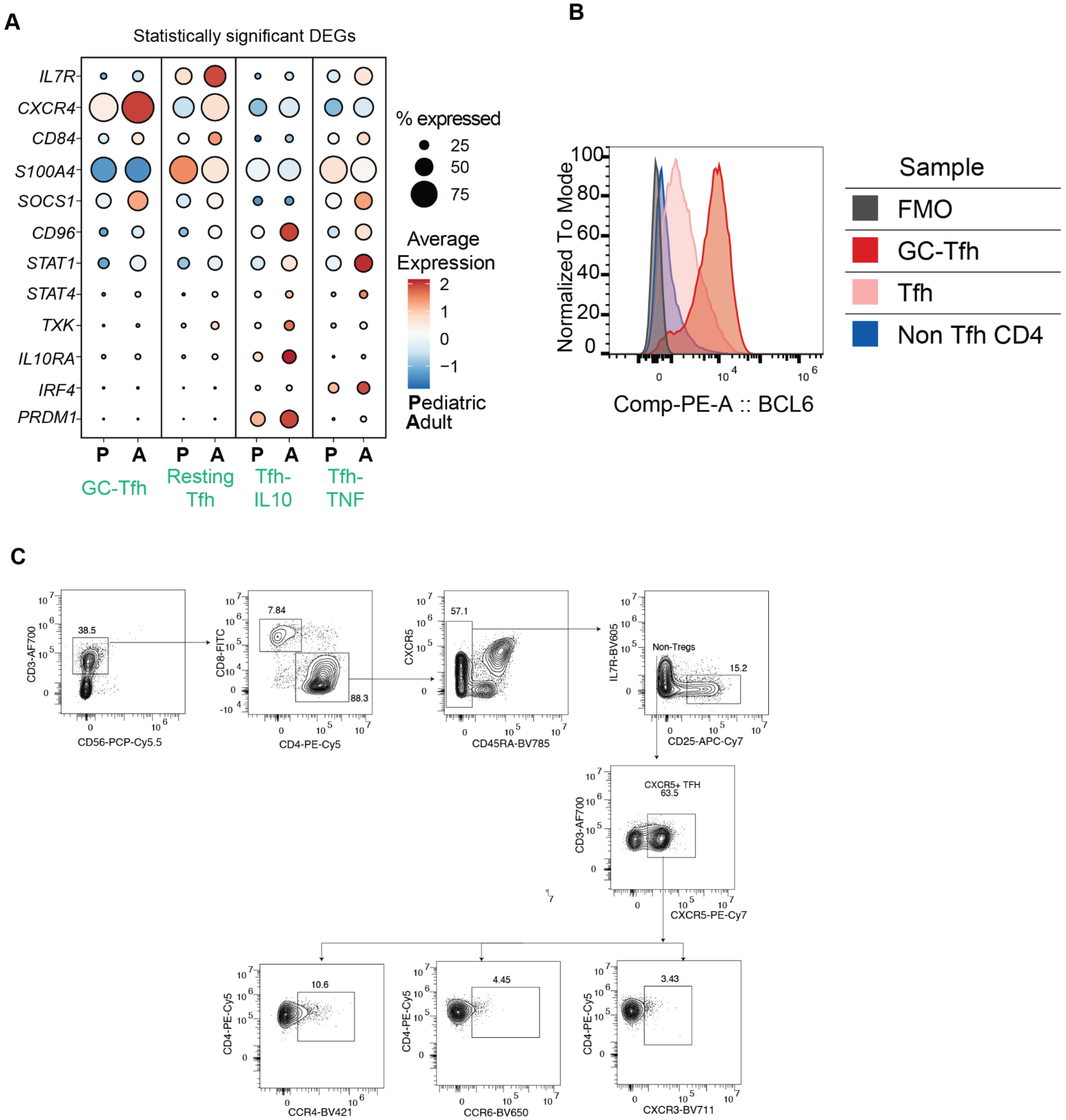


**sFig 2: Comparing Tfh phenotypes from pediatric donors and adults.**

(A) Bubble plot comparing key Tfh genes differentially expressed between pediatric donors (denoted as P) and adults (denoted as A). The size of the bubble indicates the proportion of cells within the cluster expressing the marker, and the color indicates the magnitude of expression ranging from blue (low) to red (high). (B) Representative histogram of intranuclear BCL6 expression in GC-Tfh, Tfh, and non Tfh CD4 subsets in human tonsils. (C) Gating strategy for characterization of Tfh subtypes (Tfh1, Tfh2, and Tfh17) based on chemokine receptor expression.

fig S3

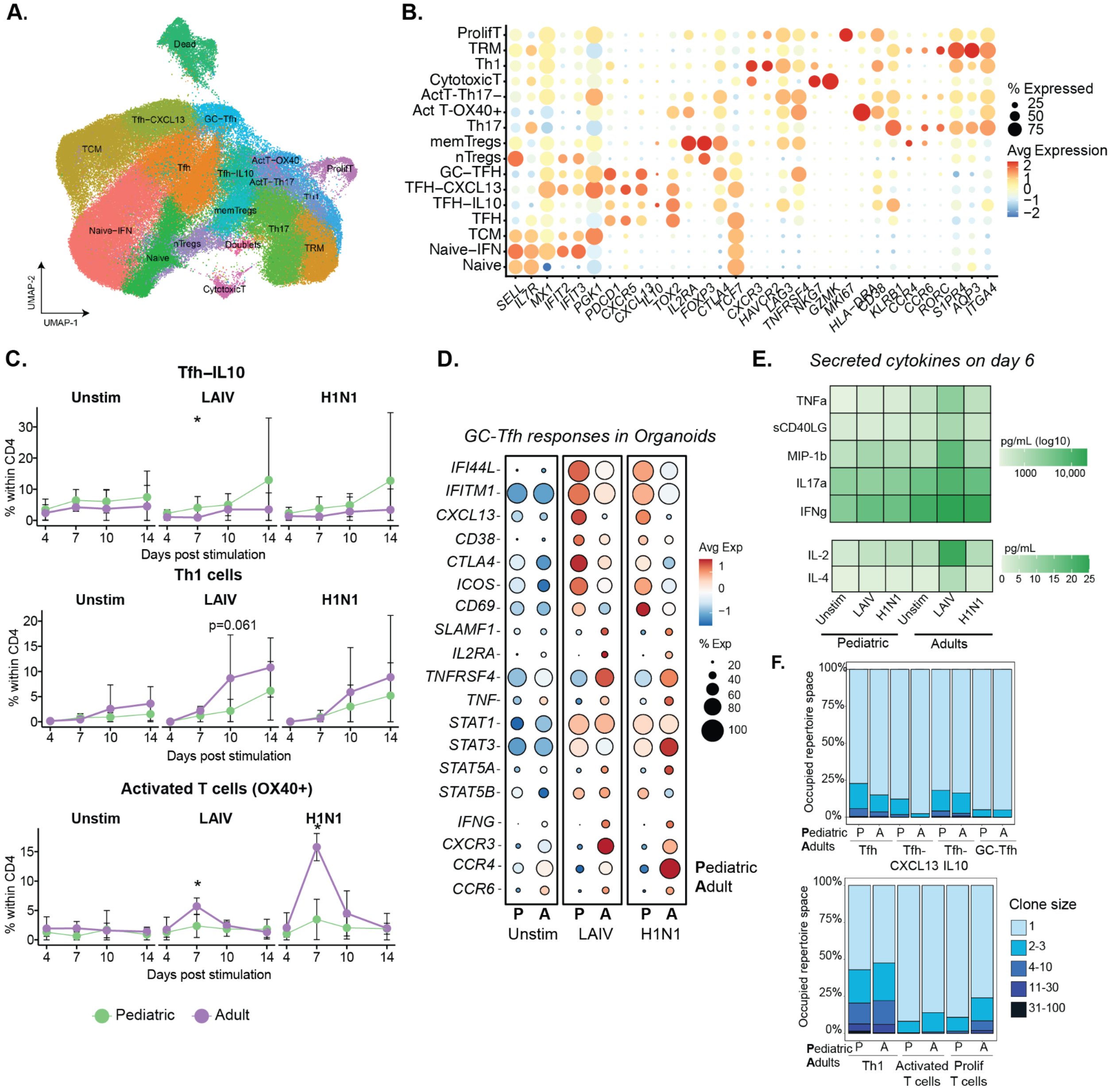

**sFig 3: scRNA-seq profiles of flu-stimulated tonsil organoids**

(A) UMAP of CD4 T cells from tonsil organoids generated from pediatric and adult donors. CD4 T cells were sorted from flu-stimulated and unstimulated organoids on days 4, 7, 10, and 14 post stimulation. (B) Bubble plot comparing gene expression markers among CD4 subsets in the tonsils. (C) Line graphs comparing proportions of T cell subsets in immune organoids over time (x-axis) within the total CD4 compartment (y-axis). At each time point, group differences were compared using unpaired Mann Whitney U test (* - p<0.05). (D) Bubble plot comparing key genes differentially expressed in GC-Tfh between pediatric donors (denoted as P) and adults (denoted as A) in immune organoids on day 7. (E) Heatmap of secreted cytokines in stimulated and unstimulated organoids from pediatric and adult donors. (F) Stacked bar graphs comparing clone size distributions among different Tfh subsets in pediatric and adult donors.

**fig S4**

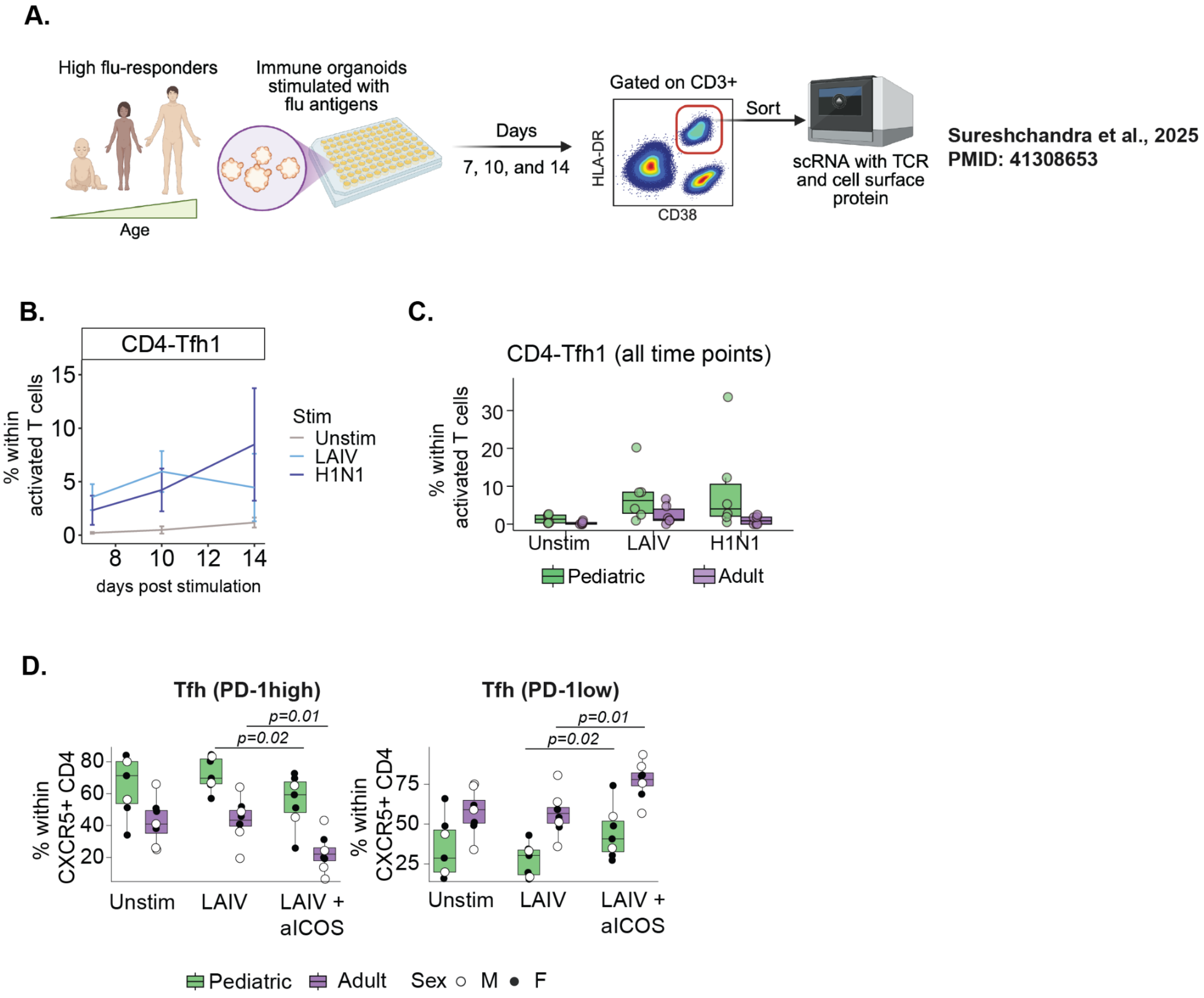


**sFig 4: Correlating Tfh responses in organoids with antibody responses.**

(A) Experimental design for deep characterization of antigen-specific Tfh responses in immune organoids (B) Line graph comparing Tfh1 frequencies in immune organoids as a percentage of activated CD4 T cells (C) Box plots comparing activated Tfh1 frequencies across stimulation conditions in pediatric and adult donors. Graph includes all time points for all donors. (D) Frequencies of PD-1 high and PD-1 low Tfh, as a percent of CXCR5+ CD4 T cells. Boxplots show the median, with hinges indicating the first and third quartiles and whiskers indicating the highest and lowest values within 1.5 the interquartile range. Within each age group, stimulation specific differences were tested using paired Wilcoxon signed-rank tests. For age-differences within each stimulation, unpaired Mann-Whitney U tests were performed.

fig S5

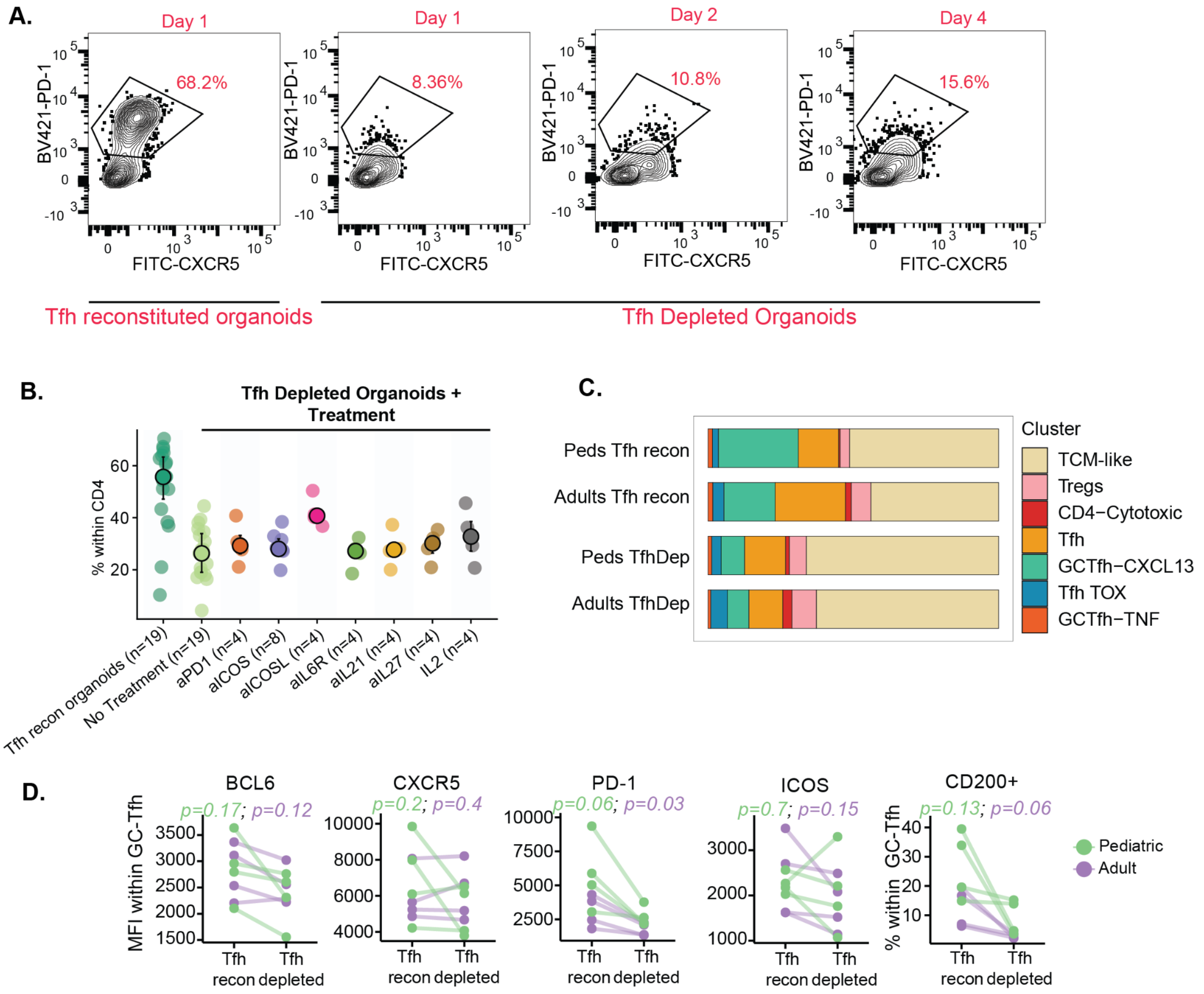


**sFig 5: Tfh depletion strategy in tonsil organoids.**

(A) Representative flow plots demonstrating the emergence of new Tfh in depleted cultures on days 1, 2 and 4 (B) Dot plots highlighting frequencies of Tfh in reconstituted and depleted cultures with or without treatment with neutralizing antibodies. X-axis denotes the neutralizing factor in each experiment relative to reconstituted cultures. (C) Stacked bar graphs comparing distribution of Tfh subsets in depleted and reconstituted cultures, aggregated across pediatric and adult donors. (D) Dotplots comparing protein expression of key Tfh markers in Tfh reconstituted and depleted cultures from pediatric and adult donors.

fig S6

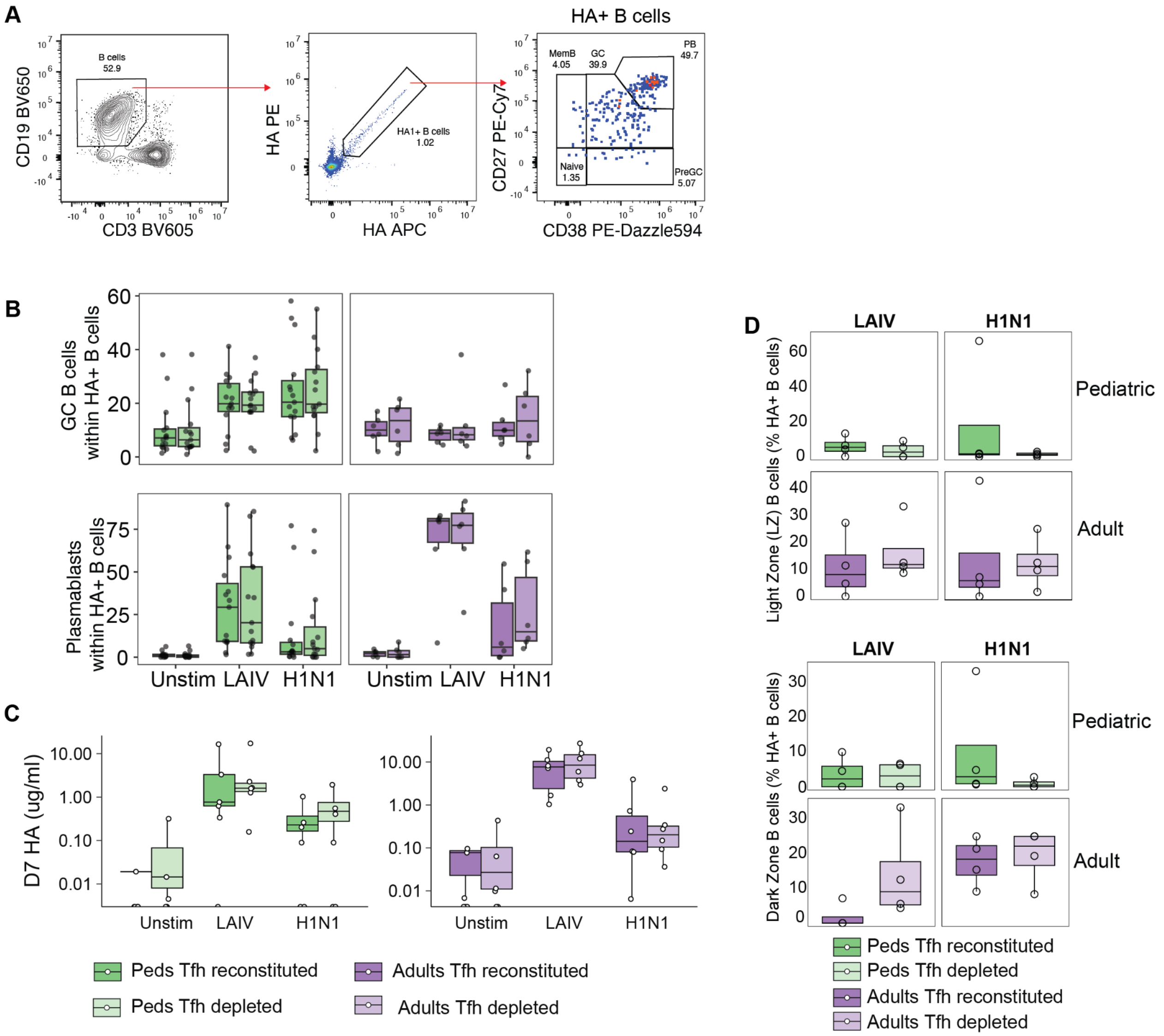


**sFig 6: Characterization of B cell responses following Tfh depletion**

(A) Gating strategy for labeling and identification of HA+ B cells (B) Boxplots comparing frequencies of HA+ cells with a GC and plasmablast phenotype in Tfh depleted and reconstituted cultures on day 7 (C) Boxplot comparing ug/mL of HA-specific antibodies in organoid supernatants on day 7 in Tfh depleted and reconstituted cultures. (D) Dotplots comparing Light Zone (top) and Dark Zone (bottom) phenotypes of HA+ cells in LAIV stimulated organoids with or without Tfh.
